# Studying macromolecular composition in cell-cell interfaces using 3D membrane reconstitution systems

**DOI:** 10.1101/2025.11.01.686034

**Authors:** Franziska Ragaller, Amelie Maribel Schneider, Ellen Sjule, Renhua Sun, Xiao Han, Edward Jenkins, Michael Dustin, Adnane Achour, Erdinc Sezgin

## Abstract

During direct communication between two cells, the plasma membranes of each cell serve as a platform for ligand-receptor interaction initiating downstream signalling cascades. In immune cell signalling, this cell-cell interface – the immune synapse – is highly spatiotemporally organized. Multiple stimulatory and co-stimulatory signals need to be integrated over time to ensure proper immune cell function. This process is still not fully understood given the vast complexity of interactions between proteins, lipids, glycocalyx and associated cortical actin cytoskeleton. To examine the impact of a single component, the use of model membrane systems has increased. Here, we developed a fully artificial system to study the interface between two vesicles and a semi-artificial one between a live cell and a vesicle to reconstitute 3D contacts. We investigated the distribution and reorganization of immune cell proteins at artificial and semi-artificial contacts. Using our vesicle-vesicle system, we show the enrichment and depletion of different proteins in the synapse. Using the cell-vesicle system we showed how different peptides with varying affinity presented by the same MHC class I affect T cell activation. We further explored the distribution of glycocalyx elements at the cell-cell contact and showed differential partitioning of different sugar moieties in the interface. While we focused on the T cell interface here, our model systems are powerful tools to study distribution and reorganization of lipids, proteins and glycocalyx components at any cell-cell contact.

## Introduction

Cell-cell communication is needed to react and adapt to environmental cues and thus crucial for the survival of multicellular organisms. To integrate external signals and mediate subsequent functions, highly sophisticated inter-and intracellular signalling networks have evolved (Nair et al., 2019). Initial events in these signalling cascades occur at the plasma membrane, which not only serves as a platform for receptors and ligands to interact, but also influences the nature and dynamics of this interaction (Grecco et al., 2011; Sezgin et al., 2017). In the context of immune cell signalling, a highly spatiotemporally organized contact is formed between different immune cells or immune cells and their target, called the immune synapse (Dustin, 2014; Monks et al., 1998). The immune synapse, best described for T cells, is spatially organized in supramolecular activation clusters, characterized by abundance or exclusion of certain proteins (Dustin, 2014). For an immune cell to decide how and when to respond, multiple stimulatory and inhibitory signals need to be integrated over time (Céspedes et al., 2021; McKeithan, 1995). To fully understand this process, the role of individual components needs to be dissected in time and space. However, this is not trivial due to the complexity at the contact, including protein-protein interactions (Dustin, 2014), lipid-protein interactions (Cho and Stahelin, 2005; Lemmon, 2008), glycocalyx involvement (Möckl, 2020), and cytoskeleton activity (Ritter et al., 2013).

By facilitating the examination of individual components involved in immune cell interactions in a live-cell context, model membrane systems of controllable complexity have gained popularity. These model membrane systems allow control over lipid composition of the membrane and proteins attached to functionalized lipids within the membrane. The use of supported lipid bilayers (SLB) on flat glass surfaces facilitated crucial advancements of our knowledge on immune cell interfaces and activation (Bertolet and Liu, 2016; Dustin et al., 2007; Grakoui et al., 1999; Johnson et al., 2000; Lopes et al., 2017). Further, SLBs allow for advanced imaging techniques such as total internal reflection fluorescence microscopy (TIRFM) and single-molecule localization microscopy (SMLM) (Brameshuber et al., 2022). However, the glass support impacts the diffusion behaviour of lipids and attached proteins (Beckers et al., 2020; Sezgin and Schwille, 2012) and increases the stiffness of the membrane beyond what immune cells typically encounter (Bufi et al., 2015; Lippert et al., 2021). Both T and B cells are highly sensitive to the stiffness of their substrate, with an increase in stiffness correlating to increased triggered signalling (Jin et al., 2019; Judokusumo et al., 2012; Lambert et al., 2017; Majedi et al., 2020; Saitakis et al., 2017; Shaheen et al., 2017; Zeng et al., 2015). Additionally, using SLBs, the cell-cell interface can only be investigated in 2D; however, cells normally form a 3D contact (Glazier and Salaita, 2017). These limitations can be overcome with the use of free-standing systems such as giant unilamellar vesicles (GUVs), allowing for reconstitution of 3D contacts closer to physiological conditions. GUVs can mimic cells well in terms of size, ranging from 1-100 μm in diameter, and also in flexibility, finite surface area, tension, deformability, and stiffness (Fenz and Sengupta, 2012; Morales-Penningston et al., 2010; Schmid et al., 2015). GUVs have previously been used to mimic immune cell interfaces in combination with themselves or SLBs (Carbone et al., 2017; Schmid et al., 2016) and in combination with cell lines (Jenkins et al., 2019).

Built on these previous reconstitution systems, we have developed two widely applicable model systems to study cell-cell interfaces. We explored the formation of completely artificial contacts using only GUVs, as well as semi-artificial contacts using GUVs in combination with T cell lines or primary human CD8^+^ T cells. We prepared GUVs by electroformation, comprising our lipids of choice and nitrilotriacetic acid (Ni-NTA)-functionalized lipids to attach His-tagged proteins to the GUV surface. We examined the distribution of adhesion molecules, T cell receptors (TCR) and peptide major histocompatibility complexes (pMHC), checkpoint inhibitors, and heavily glycosylated proteins at the artificial and semi-artificial contact. We further investigated ZAP70 recruitment and Ca^2+^ signalling as indicators for T cell activation in our semi-artificial system using both confocal and lattice light-sheet microscopy as a function of peptide affinity to TCR. Finally, we explored the glycocalyx distribution at reconstituted immune cell interfaces using lectins with specificity for different sugar residues.

While we have mainly investigated the T cell contact, the model systems presented here can be applied to a wide range of cell-cell interactions. The feasibility and versatility of these model systems make them a powerful tool to study the distribution and reorganization of lipids, proteins and glycocalyx components at 3D cell-cell interfaces.

## Methods

### Materials

We used the following lipids and fluorescent lipid probes: 1-palmitoyl-2-oleoyl-glycero-3-phosphocholine (POPC, 16:0-18:1 PC), 1-palmitoyl-2-oleoyl-sn-glycero-3-phospho-L-serine (POPS, 16:0/18:1 PS), 1,2-dioleoyl-sn-glycero-3-[(N-(5-amino-1-carboxypentyl)iminodiacetic acid)succinyl] (nickel salt) (18:1 DGS-NTA (Ni)), 23-(dipyrrometheneboron difluoride)-24-norcholesterol (TopFluor™ Cholesterol), 1-palmitoyl-2-(dipyrrometheneboron difluoride) undecanoyl-sn-glycero-3-phosphoethanolamine (TopFluor™ PE) (Avanti Polar Lipids), Abberior-STAR RED-PE (ASR-PE, Abberior GmbH) and NR12A (Danylchuk et al., 2019).

We used the following recombinant proteins obtained from Sino Biological: human CD2 (His-Tag,10982-H08H), CD58/ human LFA-3 (His-Tag, 12409-H08H), PD1/human PDCD1 (His-Tag,10377-H08H), PD-L1/human B7-H1/ CD274 (His-Tag, 10084-H08H), CD80/human B7-1 (His-Tag, 10698-H08H), human CTLA-4 (His-Tag,11159-H08H), NCR3/human NKp30 (His-Tag, 10480-H08H), human B7-H6 (His-Tag, 16140-H08H), human SLAMF6/human Ly108 (His-Tag, 11945-H08H), human CD84 (His-Tag, 10100-H08H), human SIRPαV2 (His-Tag, 30014-H08H), human CD47 (His-Tag, 12283-H08H), CD45 (ECD, His-Tag, 14197-H08H). We also used the proteins: CD43/ human leukosialin (His-Tag, CD3-H52H9, ACROBiosystems), MUC1-Alexa Fluor™ 488 (kindly provided by Carolyn Shurer), MHC class I H-2D^b^ presenting the LCMV-derived gp33 peptide (KAVYNFATM) (Achour et al., 2002; Duru et al., 2020), MHC class I (HLA-A*02:01) presenting the tumour-associated antigen NY-ESO1-9V peptide (SLLMWITQV; from now on 9V) (Chen et al., 2005), MHC class I (HLA-A*02:01) presenting the NY-ESO1-3P9V peptide variant (SLPMWITQV; from now on 3P9V), P14 TCR (Duru et al., 2020), 1G4 TCR (Chen et al., 2005), Wheat Germ Agglutinin (WGA) Alexa Fluor™ 647 Conjugate (ThermoFisherScientific), Aleuria Aurantia Lectin (AAL) (L-1390-2, Vector Labs), Sambucus Nigra-I-Agglutinin (SNA) (L-1300-5, Vector Labs), Maackia Amurensis-II Lectin (MAL-II) (L-1260-2, Vector Labs). We used Alexa Fluor™ 488 or 647 NHS Ester (Succinimidyl Ester) (ThermoFisher Scientific) for protein labelling.

NaCl, HEPES and CaCl2 were obtained from Sigma-Aldrich (St. Louis, MO, USA). Gibco™ PBS, RPMI (with L-glutamine), fetal bovine serum (FBS), Gibco™ L-glutamine (200 mM,100x), Gibco™ sodium pyruvate (100 mM), Gibco™ MEM non-essential amino acids solution (100x), Gibco™ HEPES buffer solution (1M, 100x), Gibco™ Leibovitz’s L-15 Medium (no phenol red) and Cytiva Ficoll-Paque™ PLUS Media were obtained from ThermoFisher Scientific. Recombinant human IL-2 protein was obtained from BioTechne. Fluo-4, AM was obtained from ThermoFisher Scientific.

### Protein expression & purification

To align with the experimental design, the histidine tag (His-tag) was constructed on the C-terminus of the HLA-A*02:01 heavy chain, the P14 TCR β-chain, and the 1G4 TCR β-chain, respectively. The production of both HLA-A*02:01/9V and HLA-A*02:01/3P9V as well as the H-2D^b^/gp33 was performed via refolding, following the same procedure previously described for H-2D^b^/gp33 (Achour et al., 1999; Duru et al., 2020). The P14 TCR was produced and refolded by dilution, then purified using ion exchange and size exclusion chromatography, as previously described (Duru et al., 2020). The 1G4 TCR was produced following previously published protocols (Chen et al., 2005).

### Fluorescent labelling of proteins

Proteins were fluorescently labelled using Alexa Fluor™ 488 or 647 NHS Ester (from here on AF488 or AF647). The fluorescent dyes were dissolved in DMSO to 5 mg/ml. The proteins were dissolved according to the manufacturer’s instructions in water or the respective buffer (Supplementary Table 1). To each 1 ml protein solution, 0.1 ml of 1 M NaHCO_3_ (in water, pH 8.5) was added. The volume of added fluorescent dye was calculated according to Equation 1, thereby a molecular excess of 5 was used.

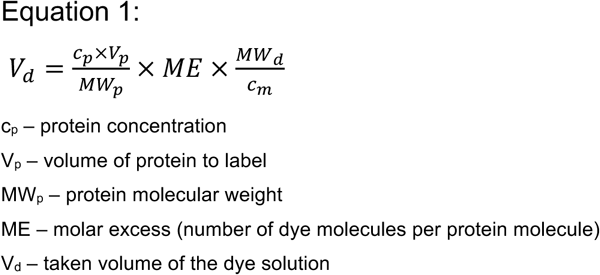

The determined fluorescent dye volume was added to the protein solution and incubated at RT for 1h under continuous rotation. For the different pMHC and TCR molecules, the reaction was carried out at 4°C overnight to prevent aggregation or denaturation of the proteins. To purify the fluorescently labelled protein, Zeba™ Spin Desalting Columns with a 7 or 40 MW cutoff (ThermoFisher Scientific) were used. To prepare the columns, they were placed in a 1.5 ml tube and centrifuged at 1500 xg for 1 min to remove the storage solution. 300 μl of respective buffer was added to the top of the resin and centrifuged at 1500 xg for 1 min. This wash step was repeated twice before the protein solution was added and again centrifuged. This protein purification process was repeated once more with a fresh column. The concentration of purified protein was determined using a Nanodrop and then aliquoted and stored at-20°C or-80°C until further use.

### Surface Plasmon Resonance (SPR) measurements

All measurements were performed on a BIAcore T200 (Cytiva) at 20°C in a buffer containing 10 mM HEPES (pH7.4), 150 mM NaCl, 0.005% Tween-20, 3 mM EDTA. Soluble 1G4-his_6_ was noncovalently captured via an anti-His antibody immobilized on a CM5 sensor chip using standard amine coupling, using HLA-A*02:01 as the mobile phase. Approximately 5000 response units (RU) of anti-His antibody were immobilized, enabling capture of ∼1000 RU of 1G4-his_6_ (TCR) (1 μM). A reference surface was prepared in parallel without 1G4-his_6_. Freshly prepared HLA-A*02:01/peptide complexes (up to 100 μM) were injected over both surfaces at a flow rate of 30 μL/min, using a series of at least ten 2-fold dilutions, with three concentrations repeated in duplicate. The sample rack was maintained at 4 °C throughout the run. Chip surfaces were regenerated with 0.1 M Glycine-HCl (pH 2.5), 500 mM NaCl, and 0.05% Tween-20 at 30 μL/min. Specific binding signals were obtained by subtracting the response from the reference surface to eliminate bulk refractive index effects and non-specific interactions. SPR data was analyzed with BIAevaluation 3.0 software (Cytiva), K_D_ values were obtained from steady–state fitting of equilibrium binding curves from at least ten sample injections.

### GUV preparation

GUVs were prepared according to a previously described protocol (Sezgin et al., 2012a). GUVs of the following lipid compositions were prepared: POPC:Ni-NTA (98:2 mol%) without fluorescent lipid, with 0.1 mol% ASR-PE and with 0.01 mol% TF-PE. The lipid mixture dissolved in chloroform (6 μl of 1 mg/ml) was homogenously distributed on two platinum electrodes in a custom-built Teflon chamber, dried under nitrogen stream, placed in 300 mM sucrose solution (370 μl) and GUVs were generated by electroformation at 2 V and 10 Hz for 1 h, followed by 2 V and 2 Hz for 30 min (Sezgin et al., 2012b).

### Cell culture maintenance

J8 Jurkat T cells, J8 Jurkat cells expressing ZAP70mNG, and P14 Jurkat T cells were cultured in fully supplemented RPMI with 10% FBS at 37°C and 5% CO_2_. The cells were split regularly according to their density. Primary human T cells were isolated from peripheral blood mononuclear cells (PBMCs). PBMCs were obtained from buffy coats of healthy human donors using density gradient centrifugation. In short, blood was diluted 1:1 with PBS and layered onto Ficoll and centrifuged at 400 xg RT for 30 min (brake off). The PBMC fraction was collected and washed with PBS three times. From PBMCs human CD8^+^ T cells were isolated using the EasySep™ Human Naïve CD8^+^ T Cell Isolation Kit II (Stemcell technologies) according to manufacturer’s instructions. The isolated cells were resuspended in fully supplemented RPMI media (10% FBS, 2 mM L-glutamine, 1 mM sodium pyruvate, 1x non-essential amino acids and 10 mM HEPES) with 50 ng/mL IL-2 and subsequently activated using Human T-Activator CD3/CD28 Dynabeads™ (ThermoFisherScientific). The beads were removed via magnetic separation after 48h of activation. T cells were further expanded while refreshing their media with new IL-2 every 48-72 h. To avoid T cell exhaustion, the T cells were not used longer than 14 days after activation.

### Flow Cytometry

J8 Jurkat cells were washed twice with PBS andstained with a viability stain (BV510 from BD Bioscience) at 1:1000 in PBS for 10 min at RT, followed by washing in staining buffer (PBS with 2% FBS). Cells were then stained with antibodies against the surface proteins CD28 (CD28-APC-H7, clone CD28.2, BD Biosciences), CTLA-4 (CTLA-4-BV421, clone BNI3, BD Horizon) and PD-1 (PD1-PE-Cy7, clone EH12.1, BD Pharmigen) 1:50 in staining buffer for 10 min at RT. After washing the cells in staining buffer, flow cytometry data were acquired using a FACS Canto II system (BD Biosciences). CompBeads (BD Biosciences) were used to perform compensation for the full antibody panel and an unstained sample was included as a control. Data analysis was performed using FlowJo 10.7.1 (FlowJo LLC).

### Cell preparation for imaging

Generally, for immune synapse reconstitution experiments, the cells were prepared immediately prior to imaging. Cells were washed twice with the respective buffer of the protein (800 xg for 1 min) and resuspended in 500 μl of buffer. For additional lectin-staining, cells were incubated with 1 μg/ml WGA-AF647 or AAL-647, or 10 μg/ml SNA-AF647 or MAL-II-AF647 at RT for 20 min, washed twice with the respective buffer (800 xg for 1 min) and resuspended in 500 μl buffer. For Ca^2+^ imaging the cells were washed twice with L-15 medium (500 xg for 3 min) and resuspended in 100 μl L-15 medium. Fluo-4, AM (1 μl) was added and incubated at 37°C for 15 min before washing the cells thrice with PBS.

### GUV labelling

For immune synapse reconstitution experiments, GUVs were decorated with different fluorescently labelled or unlabelled proteins (see Supplementary Table 1 for a complete list of proteins and their fluorescent label). To a tube containing 100μl buffer, 100 μl GUV solution was added. Depending on the experiment different amounts of protein(s) were added as indicated in Supplementary Table 2. In short, a total protein amount of approximately 2.5 pmol or 1.25 pmol was added to GUVs for artificial or semi-artificial reconstitutions, respectively. This ensured better contrast in the imaging of semi-artificial contacts. For protein exclusion titration experiments, the amount of binding protein was held constant, while the amount of excluded protein was varied. GUVs were incubated with the proteins at RT for 30 min, protected from light on a rotator set to low speed (8-10 rpm). GUVs were not washed prior to imaging. For the no-protein control, cells and GUVs were stained with 100 nM NR12A.

### Fluorescent confocal imaging & lattice light-sheet microscopy

The reconstituted immune synapses were imaged in previously blocked (3 mg/ml BSA in PBS) μ-Slides (18-well glass bottom, ibidi). To assess the contacts, 80 μl of respective buffer and 40 μl of each GUV population (artificial contacts); or 40 μl GUVs and 40 μl washed cells (semi-artificial contacts) were added into the well. In confocal imaging, artificial contacts and semi-artificial contacts with Jurkat T cells were imaged at RT, whereas semi-artificial contacts with activated human primary CD8^+^ T cells were imaged at 37°C. A Zeiss LSM980 confocal microscope with a Zeiss water immersion objective, C-Apochromat 40X/1.2 NA, was utilized. 488 nm or 639 nm lasers were used to excite the AF-488 or AF-647 conjugated proteins, ZAP70mNG or fluorescent lipid dyes. We performed fast volumetric imaging with lattice light sheet microscopy (LLSM) using the Zeiss Lattice Lightsheet 7. ZAP70mNG and Fluo-4 or AF-647 conjugated proteins were excited with 488 nm or 640 nm lasers, respectively. In lattice light-sheet imaging, semi-artificial contacts with Jurkat T cells were imaged at 37°C. Time-lapse imaging of cell-GUV contacts was performed with time intervals of 5 seconds.

### Image analysis

Confocal images and their protein signal intensities were analysed using Fiji (for a detailed description of the analysis, refer to the Supplementary Information) (Schindelin et al., 2012). In short, the regions of contact and outside the contact were defined by intensity thresholding in the respective channels and manual selection. The intensity outside the contact was normalized to 1; at artificial contacts, the outside signal is the sum of the intensities of each individual GUV. The intensity at the contact was then normalised to the outside. Values higher than 1 indicate enrichment of protein at the contact, whereas values lower than 1 indicate exclusion of the protein from the contact. To generate intensity profiles along the GUVs, the VISION software (Weber et al., 2024) employing single object detection was utilized with signal intensities normalized to the maximum signal in the profile. LLSM data were deskewed using the Zeiss software ZEN 3.11, and videos were prepared using Imaris 10.2.0.

## Results

We utilized model membranes to reconstitute cell-cell contacts bottom-up with controllable lipid and protein compositions. Here, we used GUVs containing 2 mol% Ni-NTA functionalized lipids to attach His-tagged proteins to their surface (Figure 1A) (Céspedes et al., 2021; Morales-Penningston et al., 2010). By fluorescent labelling of the proteins of interest, contact formation was examined in confocal fluorescent imaging. In the next sections, we will mainly focus on proteins involved in the formation of the T cell immune synapse.

**Figure 1.**
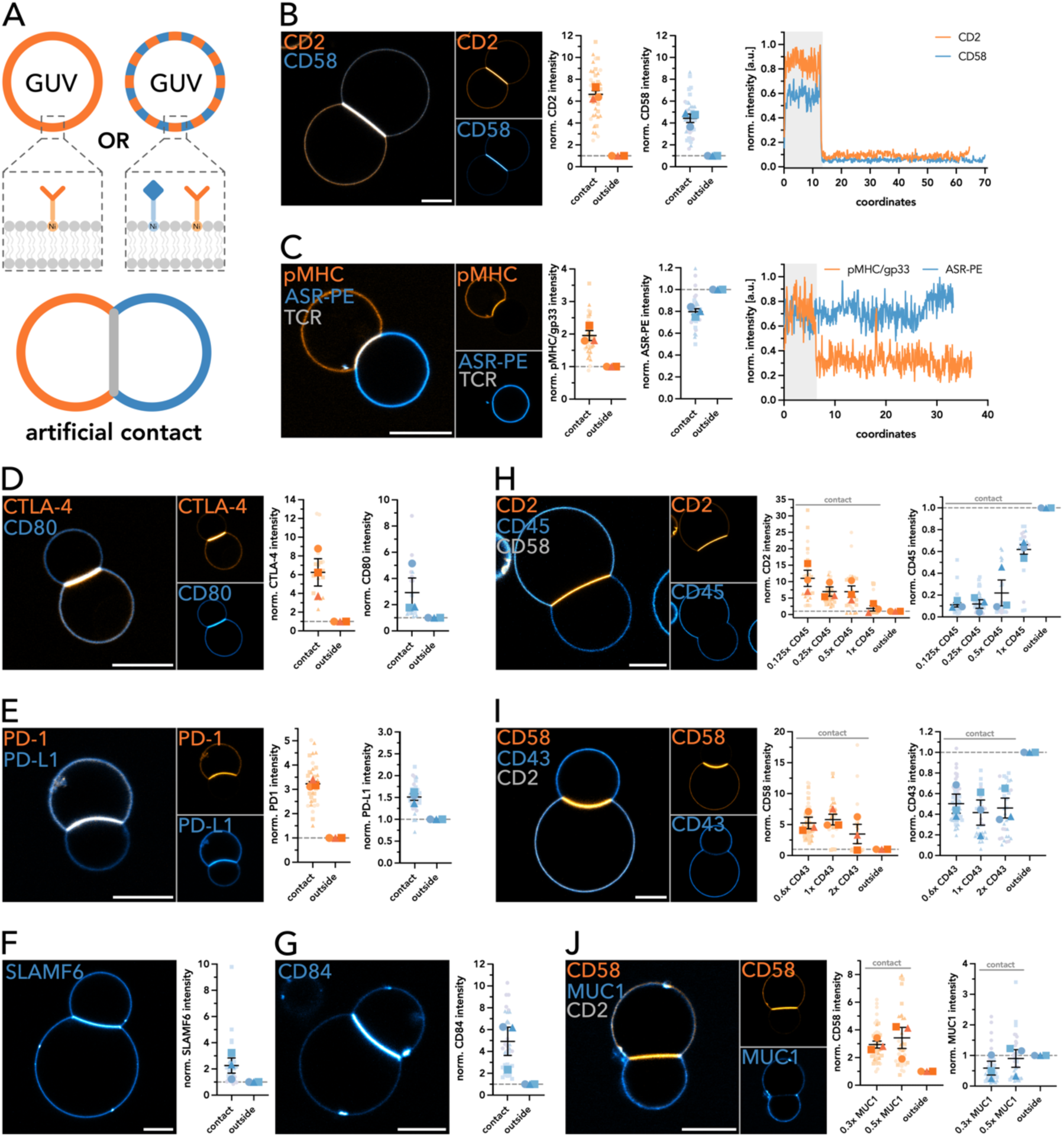
Artificial immune synapse reconstitution using free-standing GUVs. A|. Schematic representation of the artificial immune synapse reconstitution using GUVs. His-tagged fluorescently labelled proteins are attached to the GUV membranes via Ni-lipids. **B|** Representative confocal image of CD2-CD58 contact formation (left). CD2-AF488 and CD58-AF647 intensity at the contact and outside was quantified and normalized (middle). Linearized fluorescence intensity profiles of the GUVs in the confocal image with the contact highlighted in grey (right). **C|** Representative confocal image of pMHC/gp33-P14 TCR contact formation (left). P14 TCR has no fluorescent label and was reconstituted on GUVs incorporating Abberior Star Red labelled PE lipid (ASR-PE). pMHC/gp33-AF488 and ASR-PE intensity at the contact and outside was quantified and normalized (middle). Linearized fluorescence intensity profiles of the GUVs in the confocal image with the contact highlighted in grey (right). **D-E|** Representative confocal images of CTLA-4-CD80 (D) and PD-1-PD-L1 (E) contact formation (left). CTLA-4-AF488, CD80-AF647, PD-1-AF488 and PD-L1-AF647 intensity at the contact and outside was quantified and normalized (right). **F-G|** Representative confocal images of SLAMF6 (F) and CD84 (G) contact formation (left). SLAMF6-AF647 and CD84-AF647 intensity at the contact and outside was quantified and normalized (right). **H-J|** Representative confocal images of 0.125x CD45 (H), 0.6x CD43 (I) and 0.3x MUC1 (J) exclusion at the CD2-CD58 contact. Intensities of CD2-AF488 or CD58-AF647 combined with increasing amounts of CD45-AF647, CD43-AF488 or MUC1-AF488 at the contact and outside were quantified and normalized (right). Superplots show individual contacts as small symbols. Large symbols depict the mean of the individual biological replicates. Symbol corresponds to the individual biological replicate (n≥3). Standard error of the mean is shown. Scale bar corresponds to 10μm.

### Artificial model system (GUV-GUV)

#### Adhesion proteins, TCR-pMHC and checkpoint proteins enrich at the contact site

To test our artificial contact-formation system, we examined contact formation of GUVs decorated with either CD2 or CD58, adhesion proteins typically found on lymphocytes and known to bind one another (Dustin, 2014; Springer, 1990). We quantified the enrichment and exclusion of proteins at the interface (Supplementary Figure 1). The enrichment factor is the intensity fold change at the interface compared to the non-interface part (from here on called “outside”). We thereby normalized to the sum of intensities outside the contact to account for the presence of two membranes at the contact. Values higher than 1 indicate enrichment of protein at the contact, whereas values lower than 1 indicate exclusion of the protein from the contact. CD2-and CD58-GUVs formed a contact at which both CD2 and CD58 were clearly enriched compared to the outside (Figure 1B). This enrichment was also observed in the linearized profiles of the respective GUVs (Figure 1B, right panel). In contrast, neither CD58 nor CD2 alone bound themselves or were enriched at the contact (Supplementary Figure 2). We thereby confirmed that our artificial contact formation is specific to the presence of two binders on two opposite sides. We did not observe synapse formation due to unspecific binding, and lipids that are freely diffusing in the bilayer did not get enriched or depleted (Supplementary Figure 2). Besides the adhesion receptor pair CD2:CD58, we also examined contact formation of GUVs decorated with the main ligand receptor-pair involved in T cell signalling: pMHC and TCR. Here, we used the MHC class I H-2D^b^ presenting the gp33 peptide (KAVYNFATM) derived from lymphocytic choriomeningitis virus (LCMV) (from here on referred to as pMHC/gp33) and the P14 TCR, which bind each other with an affinity of 3±0.5 μM and fast kinetics with k_off_= 1 s-^1^ (Boulter et al., 2007). The P14 TCR could not be fluorescently labelled due to stability issues. GUVs reconstituted with pMHC/gp33 clearly formed contacts with P14 TCR-GUVs stained with a lipid dye, at which pMHC/gp33 enriched (Figure 1C). Interestingly, the pMHC/gp33 also formed a self-binding interface (Supplementary Figure 2).

Being the target of cancer immunotherapies, CTLA-4 and PD-1 are checkpoint inhibitor proteins expressed on the T cell surface. Binding to their ligands, such as CD80 and CD86 or PD-L1 and PD-L2, respectively, attenuates and regulates downstream T cell signalling (Krummel and Allison, 1995; Riley, 2009). GUVs decorated with CTLA-4 and CD80 formed a contact at which both proteins enriched (Figure 1D), which was not observed in the single protein controls (Supplementary Figure 2). Similarly, PD-1 and PD-L1 triggered GUV-GUV contact formation with enrichment of the proteins at the contact (Figure 1E and Supplementary Figure 2). PD1 alone did not form a real contact, whereas PD-L1 alone triggered contact formation at which PD-L1 also enriched (Supplementary Figure 2).

Inspired by this homotypic contact formation, we also tested other proteins that are reported to interact homotypically. As members of the signalling lymphocyte activation molecule (SLAM) family, SLAMF6 and CD84 share high structural similarity, are expressed by different lymphocytes, and form *trans*-homophilic interactions (Yan et al., 2007; Zhong and Veillette, 2008). Both co-stimulatory and co-inhibitory functions have been reported for SLAMF6 (Gartshteyn et al., 2022; Hajaj et al., 2020; Valdez et al., 2004) and CD84 (Cuenca et al., 2019; Yan et al., 2007). Reconstituted on GUVs, both SLAMF6 and CD84 triggered contact formation and enriched at the contact (Figure 1F-G and Supplementary Figure 2).

#### Heavily glycosylated proteins CD45, CD43 and MUC1 are excluded from the contact

At cell-cell interfaces, including immune cell contacts, numerous proteins locally enrich, whereas others get segregated or excluded locally. Segregation of CD45 has been reported from both the central and peripheral SMACs of the immune synapse (Dustin, 2014; Johnson et al., 2000), from TCR microclusters (Varma et al., 2006), at submicron-scale close contacts (Chang et al., 2016) as well as the tips of T cell microvilli (Jung et al., 2021). Therefore, we examined CD45 distribution in our artificial system. In addition to CD2 and CD58 forming a stable contact, we added increasing amounts of CD45 to both GUV species and examined contact formation (Figure 1H). CD45 was typically excluded from the CD2-CD58 contact, as clearly observable in linearized GUV profiles in Supplementary Figure 2. As our artificial system only allows passive protein distribution, driven by diffusion or mediated by protein-protein interactions, rather than by cytoskeletal or active transport mechanisms, our results suggest that the exclusion of CD45 is due to a physical mechanism, such as size-dependent segregation. The distance that CD2-CD58 proteins span at the cell-cell interface (∼130 Å) is shorter than the extracellular domain (ECD) of CD45 (even for its smallest isoform), which is rigid and usually oriented upright, leading to passive segregation of CD45 (Chang et al., 2016; Davis and van der Merwe, 1996). The ECD of CD45 is also heavily glycosylated, adding to the bulkiness of the protein (Earl and Baum, 2008). We observed that increasing amounts of CD45 hindered contact formation and decreased CD2 enrichment and CD45 exclusion at the contact (Figure 1H and Supplementary Figure 4). At the highest protein concentration, contact formation was often completely prevented (Supplementary Figure 4). Given the constant amounts of CD2 and CD58 molecules and the smaller size of the CD2-CD58 complex compared to the CD45 ECD size, the interactions between CD2 and CD58 appear to be insufficient to form a contact and exclude CD45 from the contact site, when CD45 is present in higher densities.

We also examined the distribution and influence on contact formation of the heavily glycosylated proteins CD43 and MUC1, which are highly expressed on lymphocytes and serve as a physical barrier (Ardman et al., 1992; Cyster et al., 1991; Hattrup and Gendler, 2008; van Putten and Strijbis, 2017). Similar to CD45, CD43 and MUC1 were excluded from the CD2-CD58 mediated contact and increasing glycoprotein amounts led to reduced CD58 enrichment as well as less glycoprotein exclusion (Figure 1I-J and Supplementary Figure 2). As described above, due to the large size and bulkiness of their ECDs, CD43 and MUC1 were passively segregated from the CD2-CD58 mediated contact. High concentrations of CD43 also prevented contact formation, even when the GUVs were in close proximity (Supplementary Figure 4). At the T cell immune synapse, CD43 is excluded, but according to previous studies in an active process. For example, some ezrin-radixin-moesin (ERM) family proteins link cell surface proteins to the actin cytoskeleton by interacting with the cytoplasmic tails of CD43 (Bretscher, 1999; Yonemura et al., 1998). After TCR signalling activation, CD43 is actively moved away from the immune synapse (Allenspach et al., 2001; Tong et al., 2004). Thus, CD43 exclusion from the IS seems to be both a passive process driven by kinetic segregation and an active process mediated by ERM proteins. MUC1 has also been proposed to function as a co-inhibitory molecule in T cell signalling (Agrawal et al., 2018). At high concentrations, MUC1 was also found to be enriched at parts of the GUV-GUV contact, which is counter-intuitive given its large size and extensive glycosylation. However, it was locally excluded from regions of CD58 enrichment, and vice versa (Supplementary Figure 4), indicating that at sites of CD2-CD58 binding, the opposing GUV membranes are in too close proximity for MUC1 to occupy the interface. Among the highly glycosylated proteins, MUC1 was the only one showing this behaviour at high concentrations, a phenomenon that requires further investigation.

#### The artificial model system can be used for non-T cell interfaces

The artificial GUV-GUV contact reconstitution system is also applicable to other immune cell contacts. To show this versatility, we selected two examples: *(i)* the tumour antigen B7-H6, which binds the NKp30 receptor on NK cells, triggering their activation (Brandt et al., 2009); and *(ii)* the “Don’t-eat-me” ligand CD47, expressed on all cells in the body and its inhibitory receptor SIRPα preventing uptake by phagocytes (Cockram et al., 2021; Freeman and Grinstein, 2021). B7-H6 and NKp30 (unlabelled due to instability) formed a contact at which B7-H6 was enriched, whereas in the B7-H6 only control, no protein enrichment was observed (Supplementary Figure 3). GUVs decorated with CD47 formed contacts with GUVs presenting SIRPα on their surface, resulting in CD47 enrichment at the interface, whereas SIRPα was only slightly enriched (Supplementary Figure 3). In the individual protein controls, no clear contact formation was observed (Supplementary Figure 3).

### Semi-artificial model system (cell-GUV)

#### Adhesion molecules mediate contact formation

Next, we wanted to explore whether these protein reorganizations can be seen in semi-artificial systems where a cell is combined with a GUV. We focused on T cells and investigated contact formation with both Jurkat T cells and activated human primary CD8^+^ T cells (hereafter referred to as CD8^+^ T cells). We reconstituted proteins on the GUVs and added T cells to assess contact formation and protein distribution (Figure 2A). We started with the adhesion receptor pair CD2-CD58. Jurkat T cells formed contacts with GUVs decorated with either CD2 or CD58, and both proteins enriched at the interface, with CD2 showing stronger enrichment (Figure 2B-C, Supplementary Figure 5). This indicates that although both proteins are expressed on Jurkat T cells, CD58 is expressed at higher levels. CD8^+^ T cells also formed contacts with CD58-decorated GUVs, where CD58 was enriched at the interface. In contrast, binding to CD2 only rarely led to contact formation and enrichment of CD2 at the contact (Figure 2D-E and Supplementary Figure 6). While CD58 is expressed by a broad range of both human hematopoietic and nonhematopoietic cells, including T cells, expression of CD2 is restricted mainly to T and NK cells. CD2 is the main adhesion receptor for CD8^+^ T cells, binding to its ligand CD58 on target cells or antigen-presenting cells (APCs) (Zhang et al., 2021), explaining our enrichment results. To ensure that binding was specific to the proteins reconstituted on the GUVs, we tested for the binding of Jurkat T cells and CD8+ T cells to protein-free GUVs and found that while cells were often in close proximity to the GUVs, no real contact was formed (Supplementary Figure 5 and 6). Additionally, blocking CD2 on the GUVs using a monoclonal anti-CD2 antibody prevented contact formation between Jurkat T cells and GUVs (Supplementary Figure 5). This confirmed that T cell binding to the GUVs was indeed mediated by CD2-CD58 binding.

**Figure 2:**
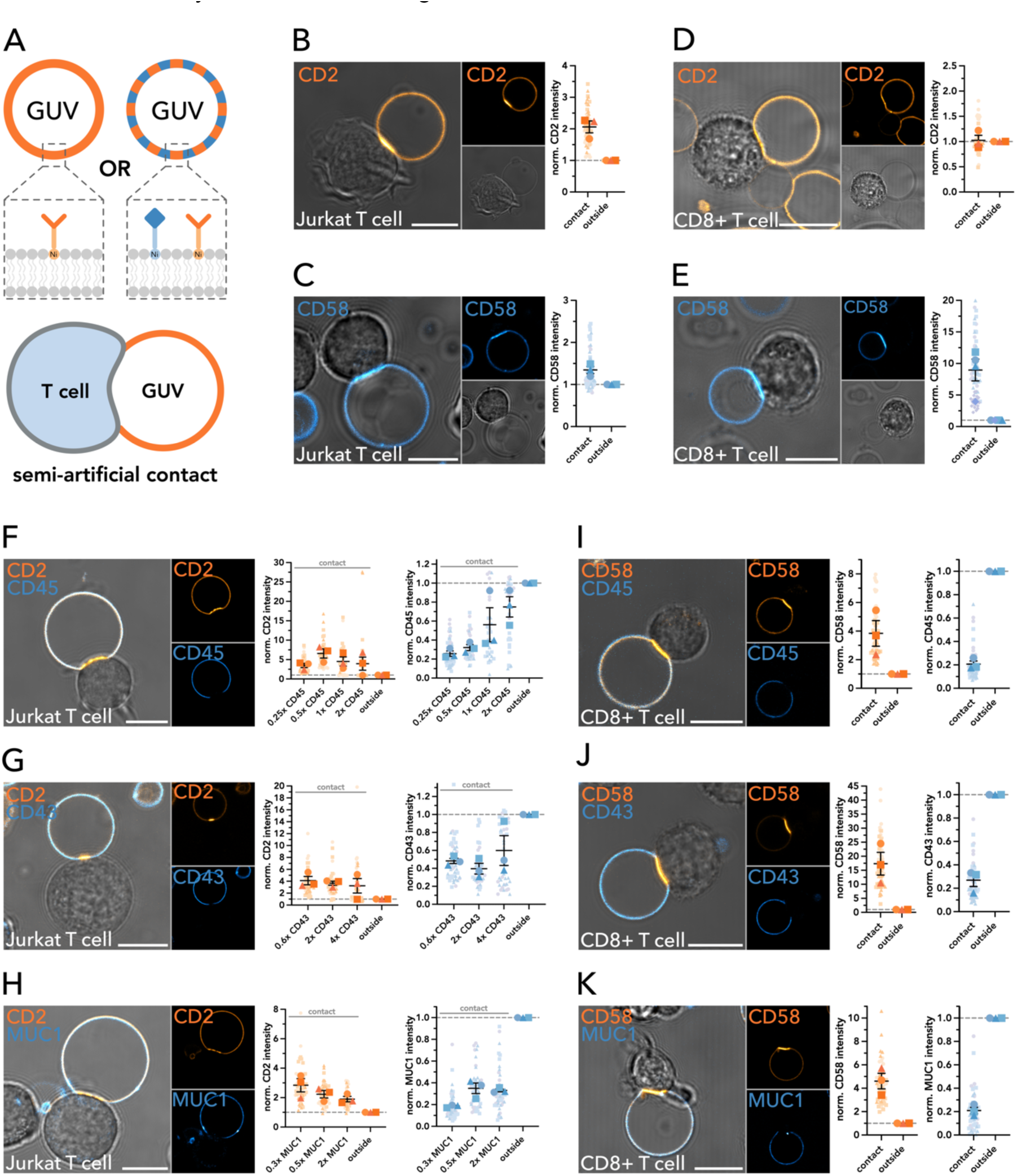
Semi-artificial immune synapse reconstitution using free-standing GUVs with Jurkat and human primary T cells. A|. Schematic representation of the semi-artificial immune synapse reconstitution using GUVs in combination with Jurkat J8 T cells or activated human primary CD8+ T cells. His-tagged fluorescently labelled proteins are attached to the GUV membranes via Ni-lipids. **B-C|** Representative confocal images of contact formation between GUVs decorated with CD2 (B) or CD58 (C) with Jurkat T cells (left). CD2-AF488 and CD58-AF647 intensity at the contact and outside was quantified and normalized (right). **D-E|** Representative confocal images of contact formation between GUVs decorated with CD2 (D) or CD58 (E) with activated human primary CD8+ T cells (left). CD2-AF488 and CD58-AF647 intensity at the contact and outside was quantified and normalized (right). **F-H|** Representative confocal images of contact formation between GUVs decorated with CD2 and 0.25x CD45 (F), 0.6x CD43 (G) or 0.3x MUC1 (H) with Jurkat T cells (left). Intensities of CD2-AF488 or CD58-AF647 combined with increasing amounts of CD45-AF647, CD43-AF488 or MUC1-AF488 at the contact and outside were quantified and normalized (right). **I-K|** Representative confocal images of contact formation between GUVs decorated with CD58 and CD45 (I), CD43 (J) or MUC1 (K) with activated human primary CD8+ T cells (left). CD58-AF488 and CD45-AF647; CD58-AF647 and CD43-AF488; or CD58-AF647 and MUC1-AF488 intensity at the contact and outside was quantified and normalized (right). Superplots show individual contacts as small symbols. Large symbols depict the mean of the individual biological replicates. Symbol corresponds to the individual biological replicate (n=3). Standard error of the mean is shown. Scale bar corresponds to 10 μm.

#### Bulky glycoproteins are effectively segregated from the contact

Similar to the artificial contacts, we also examined the distribution of glycoproteins CD45, CD43 and MUC1 at the CD2-CD58 mediated contact in the semi-artificial model system. For CD8^+^ T cells, we reconstituted CD58 on the GUV to trigger contact formation, whereas for Jurkat T cells we reconstituted CD2, as it enriched more at the GUV-Jurkat T cell contact. For both Jurkat T cells and CD8+ T cells, we observed exclusion of CD45, CD43 and MUC1 at the CD2-CD58 mediated contact (Figure 2F-K). The CD45 exclusion at T cell contacts is consistent with previous findings and has also been observed for B cell (Depoil et al., 2008), macrophage (Bakalar et al., 2018), basophil and mast cell contacts (Felce et al., 2018). With increasing amounts of CD45, CD43 or MUC1 added, we discovered a decrease in CD2 enrichment and glycoprotein exclusion at the Jurkat-T cell – GUV contact (Figure 2F-H). Interestingly, this effect was most pronounced for CD45. The enrichment of CD2 relative to the exclusion of glycoproteins was particularly evident in linearized example profiles (Supplementary Figure 5). A clear CD58 enrichment relative to the exclusion of CD45, CD43 and MUC1 was also visible at the CD8^+^ T cell - GUV contact (Figure 2I-K and Supplementary Figure 6). Compared to the fully artificial model system, the membrane distance of the cell-GUV contact is not solely defined by the CD2-CD58 complex size, and the interface is likely more heterogeneous since cell surfaces harbour many additional proteins beyond CD2 or CD58. Nonetheless, the inter-membrane distance seems small enough to enable effective size-dependent segregation of the ECDs of CD45, CD43 and MUC1.

Similar to the GUV-GUV contacts, at high glycoprotein concentrations, contact formation between Jurkat T cells and GUVs was frequently prevented, even when both were in close proximity (Supplementary Figure 7). CD43 and MUC1, among other glycoproteins, serve as a physical barrier on the cell surface and have been shown to interfere with T cell adhesion. CD43 and MUC1 are actively pushed out of the immune synapse (Allenspach et al., 2001; Tong et al., 2004); or act as coinhibitory molecule in T cell signalling, respectively (Agrawal et al., 2018; Ardman et al., 1992). Interestingly, their presence not only affects the T cell synapse, but also the phagocytic or NK cell synapse. CD43 and MUC1 are shed from the cell surface during apoptosis, thereby enhancing the uptake by phagocytic cells (Drexhage et al., 2024). In cancer, MUC1 shields adhesion and death receptors preventing immune cell binding (van Putten and Strijbis, 2017; Wi et al., 2021) and, together with CD43, increases target cell resistance to NK cell-mediated lysis (Zhang et al., 1997). As we also observed this shielding behaviour, our model systems could provide a tractable platform to study how heavily glycosylated proteins such as CD43 and MUC1 modulate or prevent specific interactions at the cell-cell interface.

#### Immune checkpoint proteins do not enrich at the contact

Further, we examined contact formation between T cells and GUVs reconstituted with immune checkpoint proteins. We chose to present CD80 on the GUV as a ligand for CTLA-4 (co-inhibitory) or CD28 (co-stimulatory) receptors expressed on T cells (Li et al., 2024), and PD-L1 as a ligand for the co-inhibitory receptor PD1 (Riley, 2009). These receptor-ligand pairs are recruited to the immune synapse (Dustin, 2014). GUVs decorated with CD80 or PD-L1 were slightly deformed upon interaction with Jurkat or CD8^+^ T cells, indicating bona fide contact formation. However, no consistent enrichment of the checkpoint proteins was observed at the contact site (Supplementary Figure 5 and 6). The Jurkat T cell clone used in this study lacks CTLA-4, and expresses only very low levels of PD1 and moderate levels of CD28, and therefore serves as a control (Supplementary Figure 5). As expected, we observed little to no PD-L1 enrichment at the Jurkat-GUV contact site, due to the low PD1 expression level. In contrast, the absence of CD80 enrichment was surprising as Jurkat cells express CD28. The reason for the missing enrichment is most likely the mechanism underlying CD28-mediated co-stimulation. CD28 binding to CD80 alone, in absence of TCR engagement, does not cause T cell activation, as it binds CD80 only monovalently and with low affinity. Upon TCR engagement, however, CD28 undergoes a conformational change that enables bivalent binding to CD80, thereby strengthening the interaction (Beyersdorf et al., 2015). As there is no TCR engagement at the Jurkat-GUV contact, CD28-CD80 interactions are likely too weak to sustain stable binding, and therefore no CD80 enrichment is observed. The lack of enrichment of CD80 and PD-L1 at the contact with CD8+ T cells was surprising, as previous studies reported upregulation of CTLA-4 and PD1 upon activation with anti-CD3/CD28 Dynabeads (Legat et al., 2013), along with sustained expression of CD28 (Li and Kurlander, 2010). The lack of additional adhesion molecules presented on the GUVs might prevent the cell from initiating contact formation, or the binding of these ligand-receptor pairs alone may be insufficient to form a stable contact, requiring additional TCR-pMHC engagement or adhesion molecule binding. In fact, a pervious study reported that proper CD28-CD80 engagement required prior adhesion mediated by CD2 (Bromley et al., 2001). Further, the upregulation of CTLA-4 or PD-1 is dynamic; hence, it is also possible that their expression at the timepoint of imaging was too low to permit detectable enrichment of the corresponding ligands presented on the GUV at the contact site. Therefore, our semi-artificial system allows to study the synergistic effect of multiple protein couples forming cell-cell contact.

#### Homophilic binding proteins rarely enrich at the contact

Reconstitution of the homophilic proteins SLAMF6 and CD84 on GUVs triggered contact formation with Jurkat and CD8^+^ T cells; however, enrichment of these proteins at the contact was only rarely observed (Supplementary Figure 5 and 6). As SLAMF6 and CD84 are expressed on activated CD8^+^ T cells (Chatterjee et al., 2011; Monel et al., 2025; Tangye et al., 2003; Yan et al., 2007) as well as on Jurkat T cells (Dragovich et al., 2019; Tangye et al., 2003), it is surprising that enrichment at the contact was only occasionally observed. The SLAM family proteins belong to the CD2 superfamily immunoglobulin (Ig) domain-containing molecules. Consequently, the homophilic interactions of CD84 and SLAMF6 share several structural and functional similarities to the CD2-CD58 interaction (Dragovich and Mor, 2018; Yan et al., 2007). The distance spanned by a CD84 *trans*-dimer (∼140 Å) falls within the range of the CD2-CD58 and TCR-pMHC interactions (Yan et al., 2007). It is therefore unlikely that additional adhesion molecules or TCR-pMHC engagement are required for proper contact formation and for CD84 or SLAMF6 enrichment. Nonetheless, the SLAMF6 or CD84 levels on the cell side may have been too low to permit detectable enrichment of the corresponding proteins presented on the GUV at the contact site.

#### Semi-artificial contact model system can distinguish peptides of different affinity

As the main ligand-receptor pair in T cell signalling, crucial for triggering downstream protein interactions and subsequent T cell activation, we examined the pMHC-TCR interaction in the semi-artificial contact model system. We decorated the GUVs either with the pMHC/gp33 alone (as described above) or in combination with CD45, and added Jurkat T cells that express the P14 TCR (Figure 3A and Supplementary Figure 8). We observed contact formation accompanied by enrichment of pMHC/gp33 at the contact site and exclusion of CD45, confirming previous literature (James and Vale, 2012). The enrichment of pMHC/gp33 was typically discontinuous, appearing as localized puncta rather than uniform accumulation along the entire contact interface. In areas with low pMHC/gp33 enrichment, CD45 was not fully excluded, a pattern that becomes especially apparent in the linearized GUV profile (Supplementary Figure 8). This suggests a more heterogeneous membrane topology, with variable distances between the cell and GUV membranes. The distance spanned by pMHC-TCR interaction (∼150 Å) is smaller than the ECD of CD45, causing its spatial segregation (Chang et al., 2016). According to the kinetic segregation model (Davis and van der Merwe, 2006), this exclusion creates a low-phosphatase environment that permits TCR phosphorylation and signalling. The size disparity is critical, as the small size of pMHC-TCR pairs promotes signalling (Choudhuri et al., 2005), whereas ECD-truncation of CD45 inhibits it (Cordoba et al., 2013). The puncta structures of pMHC/gp33 on the GUV are in accordance with the formation of TCR-pMHC microclusters during the initial stages of immune synapse formation (Bunnell et al., 2002; Campi et al., 2005). Following TCR microcluster formation, the Src family kinase Lck phosphorylates the immunoreceptor Tyrosine-based Activation Motifs (ITAMs) of the TCR, resulting in recruitment and activation of the kinase ZAP70. Activated ZAP70 then phosphorylates membrane-associated adaptor protein LAT (linker of activated T cells), which in turn recruits phospholipase Cγ (PLCγ) and promotes its activation. PLCγ activation triggers calcium release and downstream activation of Ras and PKC, ultimately resulting in T cell activation (Dustin, 2014). Therefore, recruitment of ZAP70 serves as an early indicator of T cell activation, whereas Ca^2+^ release represents a later readout. To examine ZAP70 recruitment in our semi-artificial contact model system, we used Jurkat T cells expressing ZAP70 tagged with the fluorescent protein mNeonGreen (ZAP70mNG). If T cell signalling is triggered by the proteins reconstituted on the GUV, Zap70mNG should be recruited to the contact site (Figure 3B), as demonstrated in previous studies using reconstituted T cell synapses (James and Vale, 2012; Jenkins et al., 2019). We utilized the same MHC class I molecule (HLA-A*0201) presenting two different peptides: 9V (SLLMWITQV) and 3P9V (SLPMWITQV). While the 9V variant is efficiently recognized by the 1G4 TCR (Chen et al., 2005), the 3P9V variant is not. Examined by surface plasmon resonance (SPR), the 1G4 TCR binds to the 9V and 3P9V variants with affinities of 5.2 μM and 16.6 μM, respectively (Supplementary Figure 8). We compared both enrichment at the contact and ZAP70mNG recruitment (Figure 3C). While pMHC/9V was enriched across almost the entire contact, pMHC/3P9V showed lower enrichment appearing more punctate and discontinuous (Figure 3C and Supplementary Figure 8). Consistent with the binding affinities, ZAP70mNG recruitment was higher at pMHC/9V contacts compared to pMHC/3P9V. Thus, our semi-artificial system is sensitive to differences in TCR-pMHC affinities. In pMHC-only controls, both pMHC/9V and pMHC/3P9V were enriched at the contact in the artificial system (Supplementary Figure 8). As expected, ZAP70mNG recruitment did not occur when only CD2 or CD58 were reconstituted on the GUVs (Figure 3D and Supplementary Figure 8).

**Figure 3:**
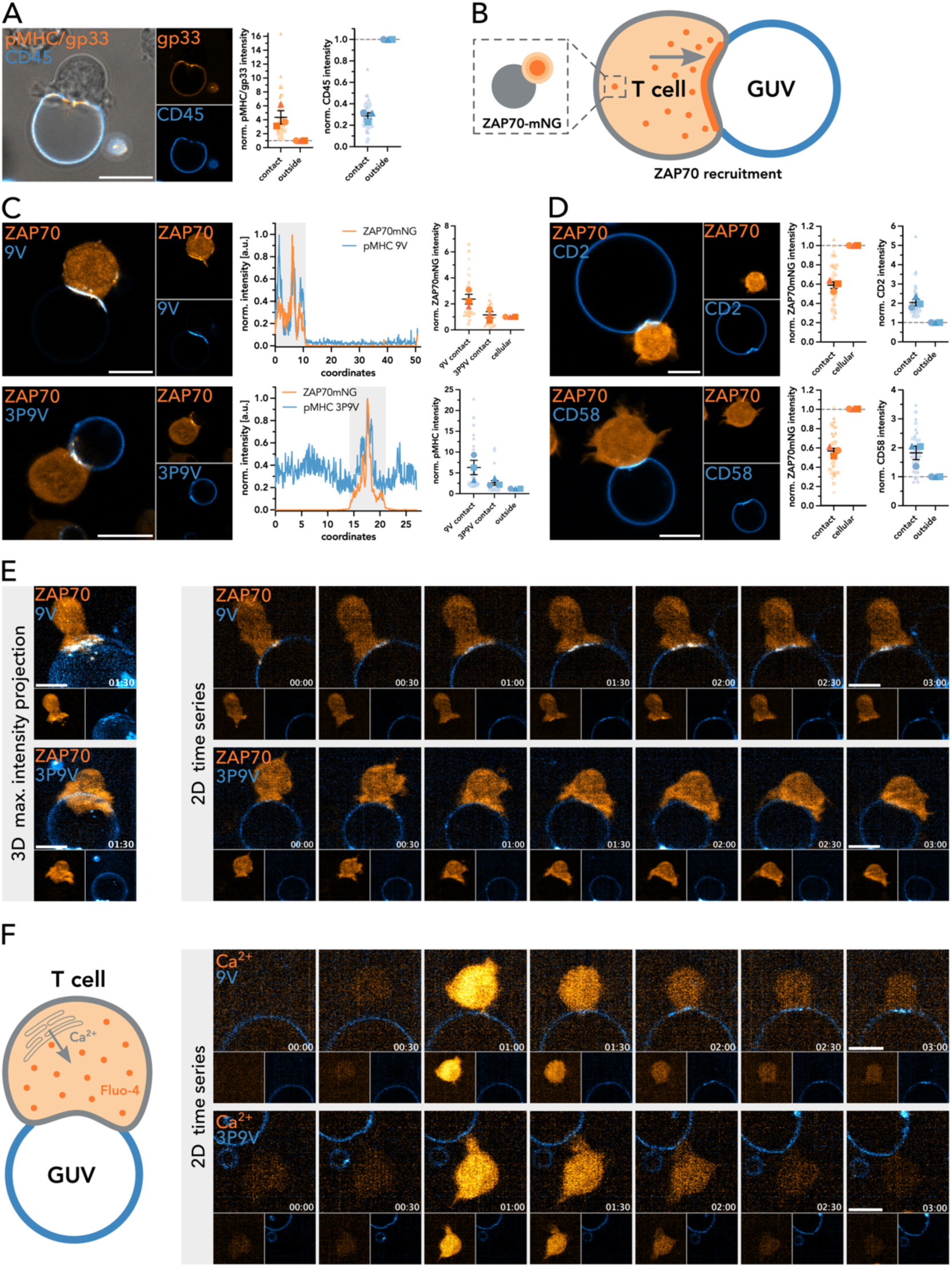
T cells get activated at TCR-pMHC mediated T cell-GUV contact. A| Representative confocal images of contact formation between GUVs decorated with pMHC/gp33 and CD45, and Jurkat P14 T cells (left). pMHC/gp33-AF488 and CD45-AF647 intensity at the contact and outside was quantified and normalized (right). **B|** Schematic representation of the ZAP70mNG Jurkat T cell line and ZAP70 recruitment to the T cell-GUV contact. **C|** Representative confocal images of contact formation between GUVs decorated with pMHC/9V or pMHC/3P9V and ZAP70mNG expressing Jurkat T cells (left). Linearized fluorescence intensity profiles of the GUVs in the confocal images with the contact highlighted in grey (middle). pMHC/9V-AF647 or pMHC/3P9V-AF647 and ZAP70mNG intensities at the contact and outside or within the cell were quantified and normalized (right). **D|** Representative confocal images of contact formation between GUVs decorated with CD2 or CD58 and ZAP70mNG expressing Jurkat T cells (left). CD2-AF647 or CD58-AF647 and ZAP70mNG intensities at the contact and outside or within the cell were quantified and normalized (right). **E|** Lattice Light Sheet Microscopy of contact formation between GUVs decorated with pMHC/9V-AF647 or pMHC/3P9V-AF647 and ZAP70mNG expressing Jurkat T cells. Maximum intensity projection of a z-stack of a cell-GUV contact at time point 1min 30s (left). Time series of the cell-GUV contact in one z-plane with every 30s shown. **F|** Schematic representation of Jurkat T cell line loaded with fluorescent dye Fluo-4 visualizing Ca^2+^ release upon cell activation due to binding to proteins reconstituted on the GUV (right). Time series of the contact between Fluo-4 loaded Jurkat T cells and GUVs decorated with pMHC/9V-AF647 or pMHC/3P9V-AF647 in one z-plane with every 30s shown. Superplots show individual contacts as small symbols. Large symbols depict the mean of the individual biological replicates. Symbol corresponds to the individual biological replicate (n=3). Standard error of the mean is shown. Scale bar corresponds to 10 μm.

These results are consistent with the role of pMHC as the primary antigenic signal in T cell activation. The CD2-CD58 interaction stabilizes contact formation and serves as a second signal in T cell activation, but does not trigger T cell signalling on its own (Binder et al., 2020; Zhang et al., 2021). The seemingly decreased ZAP70mNG intensity reflects its intracellular localization. ZAP70mNG is therefore only partly present in the contact region defined by CD58 in the analysis (see Supplementary Figure 1). To examine the distribution of pMHC/9V, pMHC/3P9V, and ZAP70 in greater detail, we performed lattice light-sheet microscopy (LLSM) for fast volumetric live-cell imaging over time with minimal phototoxicity (Figure 3E) (Chen et al., 2014). Jurkat T cells clustered pMHC/9V on the GUVs, and a local ZAP70 enrichment was observed at these pMHC/9V clusters. The ZAP70 enrichment closely tracked with the pMHC/9V clusters over time (Figure 3E). In contrast, hardly any pMHC/3P9V cluster formation or ZAP70 recruitment was observed, even though the cells actively engaged with GUVs presenting pMHC/3P9V (Figure 3E). As an additional early indicator of T cell signalling, we investigated calcium release using Fluo-4 following Jurkat T cell interaction with GUVs reconstituted with either pMHC/9V or pMHC/3P9V (Figure 3F). While calcium flashes were observed in Jurkat T cells upon interaction with GUVs for both peptides, only the pMHC/9V accumulated in clusters that moved toward the centre of the cell-GUV contact (Figure 3F and Supplementary video 5-7). Performing 4D LLSM to capture the initial contact between a cell and a GUV in the semi-artificial model system is technically challenging, as both recording the first contact event and analysing the resulting 4D data are not trivial. Nonetheless, we could showcase with limited data that live T cell contact formation and activation can be successfully examined in our model system using volumetric imaging.

#### Lectin binding suggest differential sugar residue distribution at the contact

Besides MUC1, CD43 and CD45, numerous other cell surface proteins, and a wide range of lipids, are glycosylated to varying extents. In humans, glycans are synthesized in the endoplasmic reticulum and Golgi apparatus through the stepwise addition of monosaccharides connected by glycosidic bonds (Smith and Bertozzi, 2021). The vast complexity of glycosylation arises from the ten different monosaccharides existing in either α-or β-anomers, the nature of the glycosidic bond, the various combinations of these monosaccharides into glycans of different chain lengths, and branching patterns (Möckl, 2020; Smith and Bertozzi, 2021). Glycans are attached to proteins either via the nitrogen atom of an asparagine side chain (N-glycosylation) or via the oxygen atom of threonine or serine side chains (O-glycosylation) (Möckl, 2020). The glycocalyx plays an important role for the cell as a physical protective barrier, and in cell morphology, membrane protein diffusion, cell-cell communication, immune regulation, and cancer development (Möckl, 2020). In immune cell function, glycosylation can modulate protein properties or functions, or act as a ligand for carbohydrate-specific receptors such as selectins and sialic acid-binding immunoglobulin-like lectins (siglecs) (Möckl, 2020; Smith and Bertozzi, 2021).

To further elucidate the role of the glycocalyx in immune cell contact formation, we employed lectins with defined sugar specificities in combination with our semi-artificial contact model system. Building on recent studies combining glycan microarray analysis with machine learning (Bojar et al., 2022) and super-resolution enabled cell glycotyping (Tholen et al., 2025), we examined the distribution of four lectins: (i) wheat germ agglutinin (WGA), a broadly specific lectin recognizing multiple terminal sugar residues includingGlcNAc, GalNAc and sialic acid (Neu5Ac); (ii) Aleuria aurantia lectin (AAL), which specifically recognizes α-linked fucose with various linkages; (iii) Sambucus nigra-I-agglutinin (SNA), which binds α2,6-linked sialic acid (primarily sialylated LacNAc structures); and (iv) Maackia amurensis-II lectin (MAL-II), which attaches mainly to α2,3-linkedn sialic acid (primarily Galβ1-3GalNAc in O-glycans) (Figure 4A) (Bojar et al., 2022).

**Figure 4:**
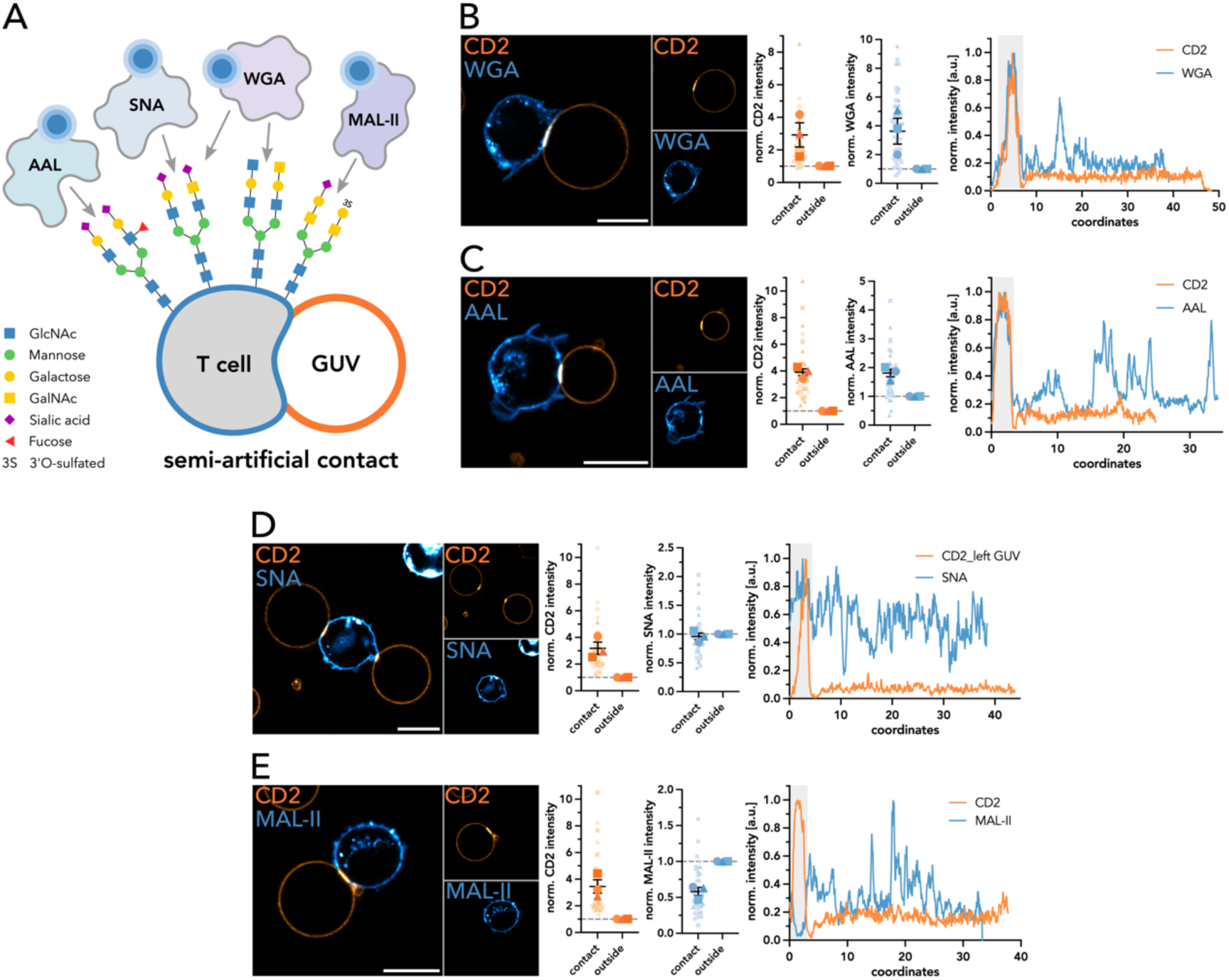
Lectins distribute differentially at the CD2-CD58 mediated T cell-GUV contact. A|. Schematic representation of the lectins used (AAL, SNA, WGA, MAL-II) to target different glycosylation patterns in the T cell glycocalyx and their distribution at the T cell GUV contact. **B-E|** Representative confocal image of contact formation between GUVs decorated with CD2 with Jurkat T cells labelled with WGA (B), AAL (C), SNA (D) and MAL-II (E) (left). CD2-AF488 intensity and WGA-AF647, AAL-AF647, SNA-AF647 or MAL-II-AF647 intensities at the contact and outside were quantified and normalized (middle). Linearized fluorescence intensity profiles of the GUV and cell stained with the respective lectin in the confocal image with the contact highlighted in grey (right). Superplots show individual contacts as small symbols. Large symbols depict the mean of the individual biological replicates. Symbol corresponds to the individual biological replicate (n=3). Standard error of the mean is shown. Scale bar corresponds to 10 μm.

Generally, lectin staining resulted in a relatively homogenous cell surface staining, interspersed with occasional larger protein clusters. Some degree of internalization was observed, especially during longer imaging periods (Figure 4B-E and Supplementary Figure 9). In Jurkat T cells, both WGA and AAL were overall enriched at the CD2-CD58 mediated contact (Figure 4B, C). SNA was neither fully enriched nor excluded from the contact with the CD2 reconstituted GUV (Figure 4D). MAL-II was excluded from the CD2-CD58-mediated contact (Figure 4E). None of the lectins bound to protein-free GUVs containing Ni-lipid, although MAL-II seldom accumulated at GUV-GUV contacts (Supplementary Figure 9). Lectin-stained cells positioned in close proximity to protein-free GUVs containing the fluorescent membrane dye Topfluore labelled PE lipid (TF-PE) did not show comparable enrichment and exclusion patterns as at the CD2-CD58 mediated contact (Supplementary Figure 9), confirming that the observed enrichment at the CD2-CD58 mediated contact was specific. Of note, the lectins used differ in valency, with WGA, AAL and MAL-II being dimers (B S and Surolia, 2017; Olausson et al., 2011; Portillo-Téllez et al., 2011) and SNA being a tetramer (Shibuya et al., 1987). Valency influences binding affinity (Riera et al., 2021), which may potentially affect lectin staining, accumulation and quantification.

Our results suggest an accumulation of GlcNAc, GalNAc (WGA), and α-linked fucose (AAL) residues at the contact, whereas α2,6-linked sialic acid neither enriches nor is excluded (SNA), and 2,3-linked sialic acid is excluded (MAL-II). Notably, CD2 contains only a single N-linked glycosylation site, predominantly with high mannose content, which is required to stabilize the domain and ensure proper binding to CD58 (Recny et al., 1992). As CD2 contains only a single glycosylation site and no lectin binding was observed on the GUV, we can exclude that the accumulation of WGA or AAL is due to CD2 enrichment. However, CD58 on the Jurkat cell side carries six N-linked glycosylation sites and has been described to be 44-68% carbohydrate (Wallner et al., 1987). Therefore, it is difficult to determine whether the observed accumulation or exclusion of specific sugar residues only reflects CD58-specific glycosylation patterns or also includes contributions from other glycoproteins or glycolipids present at the contact. Nonetheless, the strong exclusion of MAL-II suggests a more general exclusion of α2,3-linked sialic acid residues, as it is unlikely that CD58 is the only glycoprotein present at the contact. The observed enrichment of fucose residues at the contact could also reflect the accumulation of TCRs at the contact site. TCRs are highly core-fucosylated, a modification known to be important for T cell activation (Fujii et al., 2016; Liang et al., 2018).

Although further investigations employing click chemistry for specific sugar labelling (Jewett et al., 2010), advanced microscopy techniques such as Lectin-PAINT (Tholen et al., 2025), or omics approaches like glycoproteomics (Woo et al., 2017) are needed to fully elucidate the role of the glycocalyx at immune cell contact sites, we demonstrate that our system is suitable not only for studying the protein components of contact formation, but also to investigate the influence of the glycocalyx.

## Conclusion

The mechanisms governing cell-cell interfaces have long been and continue to be, a central focus in biology. Thereby, especially our understanding of immune cell contacts has greatly benefitted from the use of model membrane systems. The reduced complexity such systems enables the study of individual components from lipids and proteins to glycocalyx, within a live-cell context. Here, we established both fully artificial and semi-artificial model systems based on GUVs to mimic freestanding 3D cell-cell interfaces. While we primarily explored T cell contacts, the system can be easily adapted to investigate similarities of differences of immune cell contacts formed by other lymphocytes or phagocytes. It may also be applied to investigate differences between immune cell subsets, for example T memory versus T regulatory cells, or across cell lines, as for example Jurkat T cell immune synapses differ in topology, actin architecture, and adhesion molecule expression (Cassioli et al., 2021; Kumari et al., 2019). Importantly, this system is not limited to immune cell contacts and could be applied to mimic any type of cell-cell interface.

Mechanistically, we have shown that the semi-artificial model system can distinguish peptides of different affinity presented by the same MHC class I. It would be of interest to apply this system to a broader array of peptides, including both low-and high-affinity epitopes, to investigate their potential to induce agonist or non-agonist interactions with the TCR. The model systems presented here could also be employed to investigate *cis* protein interactions and their potential impact on *trans* protein interactions. For example, it was reported that PD-L1 and CD80 dimerize in *cis*, which inhibits *trans* CD80:CTLA-4 and PD-L1:PD1 interactions while still maintaining *trans* CD80:CD28 engagement (Zhao et al., 2019). Beyond protein distribution, we have briefly explored glycocalyx distribution at Jurkat T cell contacts using lectins. Often underestimated, the glycocalyx has immunomodulatory functions and plays an important role in cancer immune evasion (Möckl, 2020). While we cannot yet draw definitive conclusions regarding the enrichment or exclusion of specific sugar residues, the results highlight both the knowledge gap regarding glycocalyx influence at immune cell contacts, as well as the wide applicability of these model systems. Further, these model systems can be used to investigate the influence of lipids on contact formation and protein distribution. Here, we have employed simple POPC membranes; however, examination of lipid compositions of varying membrane fluidity or with the ability to phase-separate is of interest, as immune cell surface proteins have been shown to preferentially partition into distinct lipid environments at the contact interface (Shelby et al., 2023; Urbančič et al., 2021). Measuring membrane fluidity using environment-sensitive probes (Carravilla et al., 2025) at the semi-artificial contact could further elucidate the role of lipids at cell-cell interfaces. Of note, decreasing membrane fluidity following granzyme and perforin release at the immune synapse has been proposed as a protective mechanism for T and NK cells, preventing accidental death (Li and Orange, 2021; Rudd-Schmidt et al., 2019). Similarly, exposure of negatively charged phosphatidylserine on the outer T cell leaflet sequesters and inactivates perforin (Rudd-Schmidt et al., 2019). This protection mechanism could be studied in more detail using our model systems. The diffusion of lipids, proteins and glycocalyx components impacts their distribution and signalling at cell-cell interfaces. A major advantage of the freestanding model systems is that macromolecules are free to diffuse, allowing their dynamics to be quantified, using fluorescence (cross-) correlation spectroscopy.

While these model systems offer many advantages, they also have certain limitations. GUVs do not reflect the full complexity of a cell in terms of lipid-, protein-and glycocalyx diversity and lack an associated cytoskeleton. Although ECD size heavily influences diffusion behaviour and partitioning into different lipid domains (Gurdap et al., 2022), full-length proteins may behave differently, potentially experiencing steric confinement that affects their distribution at the cell-cell interface. This limitation could be overcome by functionalizing GUVs with full-length transmembrane proteins (Dezi et al., 2013). Further, imaging of the free-standing reconstituted cell contacts cannot be performed using TIRF, the method of choice for 2D SLB-cell interfaces. Nonetheless, fast volumetric imaging using LLSM of the freestanding contact is feasible, enabling the visualization of cell protrusions and their pushing and pulling of the GUV membranes.

Taken together, we established two widely applicable cell-cell interface reconstitution model systems based on GUVs (artificial or semi-artificial), which form freestanding 3D contacts. These systems are highly versatile, enabling the study of any cell-cell interface in the context of lipids, proteins, and glycocalyx components.

## Acknowledgement

We thank the SciLifeLab Advanced Light Microscopy facility and National Microscopy Infrastructure (VR-RFI 2016-00968) for their support on imaging. We would like to thank Carolyn Shurer for kindly providing a protein used in this study (MUC1-Alexa Fluor™ 488). We have been supported by Swedish Research Council Grants (grant no. 2020-02682, 2024-02993 and 2024-00289), Wellcome Leap’s Dynamic Resilience Program (jointly funded by Temasek Trust), Karolinska Institutet (2024-03250; 2024-03341; 2022-00803; 2020-00997), Cancer Research KI (2024-03488), Human Frontier Science Program (RGP0025/2022), Longevity Impetus Grant from Norn Group, Hevolution Foundation and Rosenkranz Foundation. ES is an EMBO Young Investigator (YIP-2025). The Achour research group received financial support from Vetenskapsrådet (2021-05061), Cancerfonden (24 3775 Pj), Radiumhemmets Forskningsfonder (244092) and Insamlingsstiftelsen Cancer-och Allergifonden (244092).

## Notes

### Competing Interest Statement

The authors have declared no competing interest.

