## Supplementary Information for "Studying macromolecular composition in cell-cell interfaces using 3D membrane reconstitution systems"

**Supplement Table 1: Proteins used with their respective fluorescent label and buffer.**

| Protein | fluorescent label | buffer |
|---|---|---|
| CD2 | none | PBS |
| | Alexa Fluor 488 | |
| | Alexa Fluor 647 | |
| CD58 | none | |
| | Alexa Fluor 488 | |
| | Alexa Fluor 647 | |
| pMHC/gp33 | Alexa Fluor 488 | |
| P14 TCR | none | |
| pMHC/9V | Alexa Fluor 488 | |
| pMHC/3P9V | Alexa Fluor 488 | |
| 1G4 TCR | none | |
| PD-1 | Alexa Fluor 488 | |
| PD-L1 | Alexa Fluor 647 | |
| CTLA-4 | Alexa Fluor 488 | |
| CD80 | Alexa Fluor 647 | |
| CD84 | Alexa Fluor 647 | |
| SLAM F6 | Alexa Fluor 647 | |
| Nkp30/NCR3 | Alexa Fluor 488 | |
| B7-H6 | Alexa Fluor 647 | |
| SIRP$\alpha$ | Alexa Fluor 647 | |
| CD45 (non-AB) | Alexa Fluor 647 | |
| CD43 | Alexa Fluor 488 | |
| MUC1 | Alexa Fluor 488 | |
| WGA | Alexa Fluor 647 | HBSS |
| AAL | Alexa Fluor 647 | 10 mM HEPES<br>150 mM NaCl<br>0.1 mM $CaCl_2$ |
| SNA | Alexa Fluor 647 | |
| MAL-II | Alexa Fluor 647 | |

**Supplement Table 2: Protein amounts used in different model systems.**

| | **artificial contact (GUV-GUV)** | **semi-artificial contact (GUV-cell)** |
|---|---|---|
| **single protein** | ~2.5 pmol | ~ 1.25 pmol |
| **dual protein** | | |
| contact forming protein | ~1.25 pmol | ~0.63 pmol |
| potentially excluded protein | 0.16 - 2.5 pmol | 0.16 - 2.5 pmol |

*Artificial GUV-GUV contacts – Quantification*

Image analysis and quantification was performed in Fiji (Schindelin et al., 2012) and is illustrated in Supplement Figure 1A. The original multi-channel image is split into the different channels. Thresholding on fluorescent channels was used to define areas of contact and individual GUVs. These areas were saved as ROIs (regions of interest). Additional manual selection of contact, GUV1 and GUV2 was performed ,and these regions were combined with ROIs obtained from thresholding using the "AND" function. This excludes intensity from potential bright aggregates or other GUVs not involved in contact formation in the image. The fluorescent intensity was measured for each region and background in each channel. After background subtraction the fluorescence intensity of the contact was normalized to the sum of intensities outside of the contact of GUV1 and GUV2 for each channel. The sum of intensities of the GUVs was set to 1. Therefore, normalized intensity values above 1 indicate enrichment, below 1 exclusion and around 1 neither enrichment nor exclusion from the contact. To ensure this image analysis and quantification is correct, we used GUVs incorporating the lipid dye TF-chol and added either CD58-AF647 or unlabelled CD2. CD2-CD58-mediated contact formation occurs as indicated by the enrichment of CD58 at the contact, whereas the values for TF-chol fluctuate around 1, verifying our analysis strategy (Supplement Figure 1B). The presented image analysis pipeline was followed and adapted for the semi-artificial system with unlabelled or Lectin-labelled cells. For cells expressing ZAP70mNG, the ZAP70mNG signal recruited to the contact site was normalized to the average overall intensity of ZAP70mNG in the cell.

*Artificial GUV-GUV contacts – further controls*

To ensure the artificial contact-formation system works properly, enriched signal and contact formation are indeed mediated by protein interactions and correct analysis is conducted, we performed a number of controls. The fluorescent signal observed on the GUVs stems from fluorescently labelled protein and not free dye as the dyes alone do not bind to the GUVs containing Ni-functionalized lipids (Supplement Figure 2A). Protein-less GUVs incorporating either TF-PE or ASR-PE as fluorescent lipid dyes sometimes form "contacts", however this is rare and caused by chance (Supplement Figure 2B). As the proteins get into close physical proximity upon binding and accumulating at the contact, we performed a FRET control and quantified the signal at the contact upon 488 nm excitation only in both the green (495-600 nm) and red channel (645-750 nm) and no FRET could be observed (Supplement Figure 2C). Of note, CD2-AF488 can be detected on primarily CD58-AF647 positive GUVs and vice versa (Supplement Figure 1C), which is also observed for other protein combinations. This is due to the nature of the protein reconstitution on GUVs in our system. Ni-lipids and His-tagged proteins do not from a covalent bond, but a chelation bond, so binding is reversible. As soon as the two GUV populations decorated with different proteins A and B are added together, there is an exchange of the proteins between the GUV populations (Raghunath and Dyer, 2019). This is determined by the number of Ni-lipids on the GUV, the length of the His-tag of the protein and its position in respect to the protein, and the amount of protein bound to the surface (protein crowding) (Raghunath and Dyer, 2019). Also, the protein enrichment fluctuates due to heterogeneity in lipid distribution – some GUVs will have more or less than 2% Ni-lipid in their outer lipid layer, even though all GUVs were produced from the same lipid mixture. Further, the method of protein labelling cannot ensure only one fluorescent label per protein molecule, further contributing to different degrees of enrichment of 2 proteins at the same contact.

### Supplementary Figures

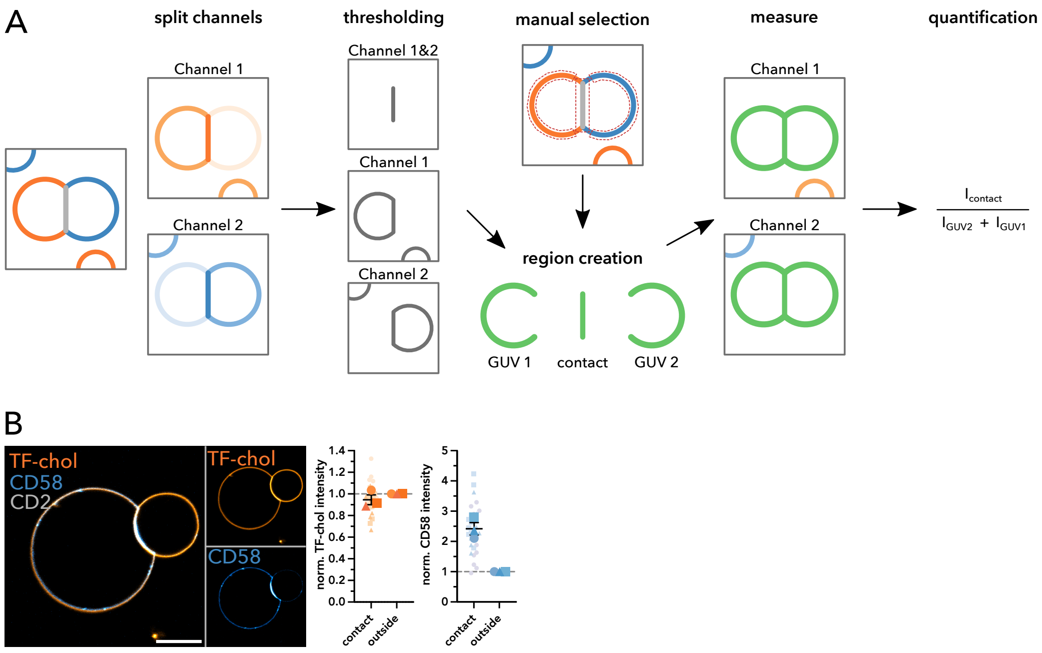


**Supplement Figure 1: Fluorescent intensity quantification.** A| Schematic representation of the image analysis using the artificial GUV-GUV system as an example. In Fiji, channels of the original image are split and the fluorescence channels are used for thresholding of the contact site and the individual GUVs. Additional manual selection of the areas is performed to ensure proper selection. The thresholded areas and manually selected areas are combined to create the regions: contact, GUV 1 and GUV 2. Fluorescence intensity in these regions is measured in each channel. After background subtraction the intensities are normalized and quantified. B| Quantification control. Representative confocal image of GUVs comprising Ni-lipid and 0.1% TF-chol decorated with either unlabeled CD2 or CD58-AF647 (left). TF-chol and CD58-AF647 intensity at the contact and outside was quantified and normalized (right).

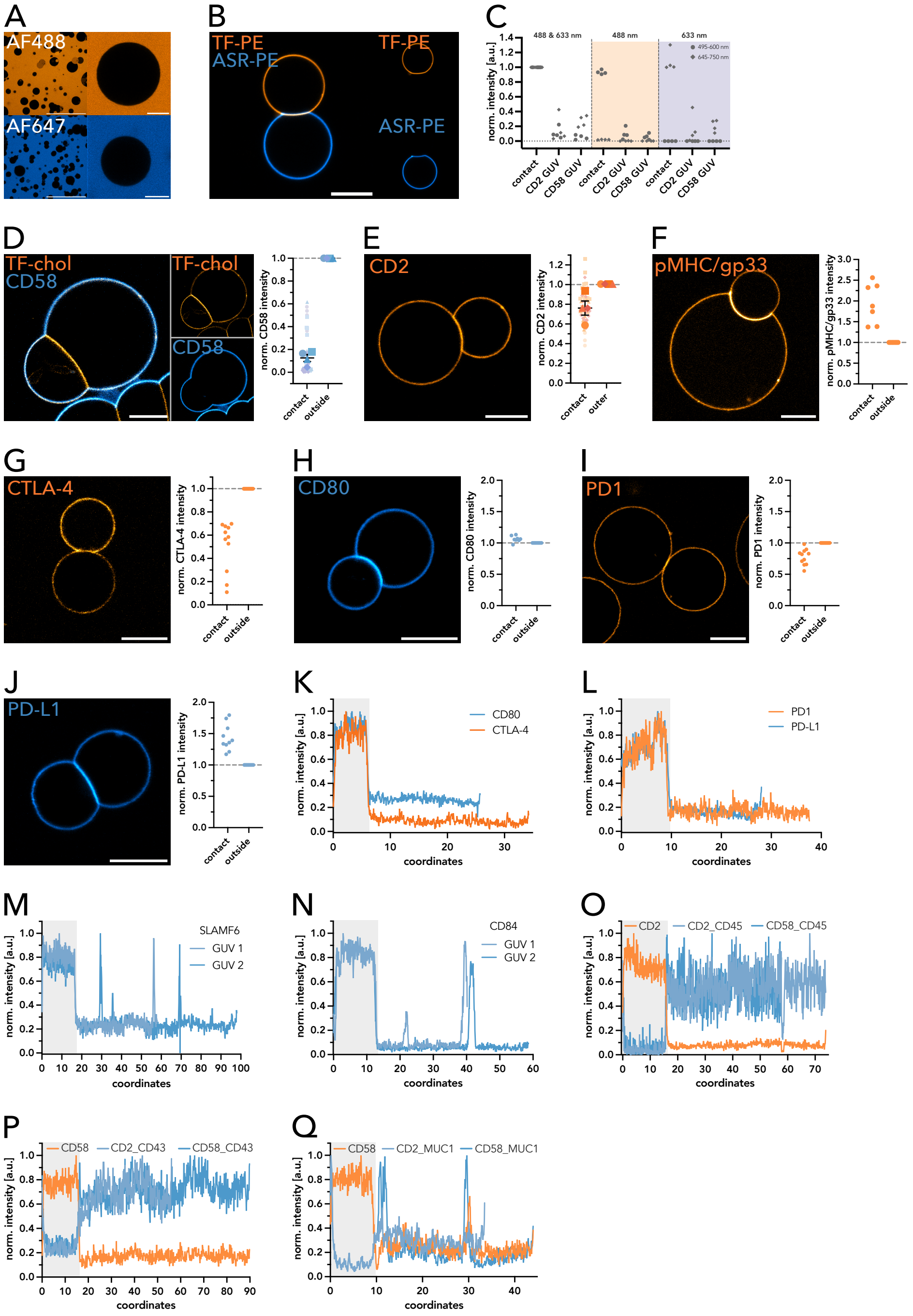


**Supplement Figure 2:** A| Alexa Fluor 488 and Alexa Fluor 647 dye control. Confocal images of GUVs comprising Ni-lipid and added fluorescent dye (1:10000). B| Contact formation control. Representative confocal image of GUVs comprising Ni-lipid and fluorescently labelled lipids (0.1% ASR-PE or 0.01% TF-PE). C| FRET control. Depicted are normalized intensities of CD2-AF488 and CD58-AF647 at the contact and outside collected in two channels (dot: 495-600 nm; diamond: 645-750 nm) with both excitation lasers, only 488 nm excitation (light orange) and only 633 nm excitation (light blue). D| CD58-AF647 only control. Representative confocal image of GUVs comprising Ni-lipid and 0.1% TF-chol decorated with CD58-AF647 (left). CD58-AF647 intensity at the potential contact and outside was quantified and normalized (right). E-J| Single protein controls. Representative confocal images of GUVs comprising Ni-lipid and with CD2-AF488 (E), pMHC/gp33-AF488 (F), CTLA-4-AF88 (G), CD80-AF647 (H), PD1-AF488 (I), PD-L1-AF647 (J) (left). Fluorescent intensity at the potential contact and outside was quantified and normalized (right). K-L| Linearized fluorescence intensity profiles of the GUVs decorated with either CTLA-4-AF488 and CD80-AF647 (K) or PD-1-AF488 and PD-L1-AF647 (L) with the contact highlighted in grey in the confocal images of Figure 1 D-E. M-N| Linearized fluorescence intensity profiles of the GUVs decorated with either and SLAMF6-AF647 (M) or CD84-AF647 (N) with the contact highlighted in grey in the confocal images of Figure 1 F-G. O-Q| Linearized fluorescence intensity profiles of the GUVs decorated with CD2 and CD58 and additional CD45-AF647 (O), CD43-AF488 (P) or MUC1-AF488 (Q) with the contact highlighted in grey in the confocal images of Figure 1 H-J. Superplots show individual contacts as small symbols. Large symbols depict the mean of the individual biological replicates. Symbol corresponds to the individual biological replicate ($n \geq 3$). Standard error of the mean is shown. Scale bar corresponds to 10 $\mu$m (small field of view; single GUVs) or to 100 $\mu$m (large field of view; A).

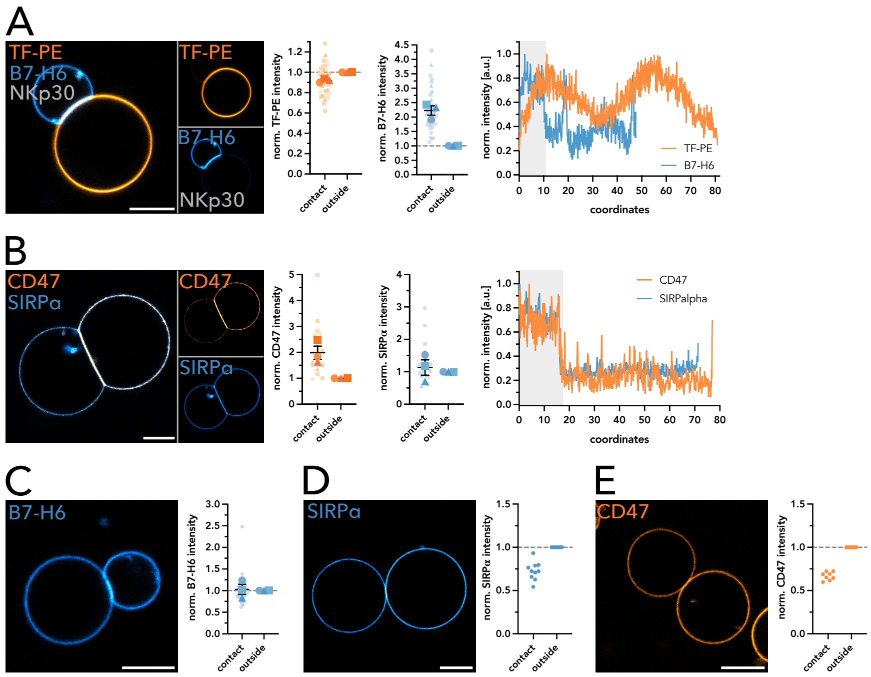


**Supplement Figure 3:** A| Representative confocal image of Nkp30-B7H6 contact formation (left). Nkp30 has no fluorescent label and was reconstituted on GUV incorporating TF-PE. TF-PE intensity and B7H6-AF647 at the contact and outside was quantified and normalized (middle). Linearized fluorescence intensity profiles of the GUVs with the contact highlighted in grey in the confocal image (right). B| Representative confocal image of CD47-SIRPα contact formation (left). CD47-AF488 and SIRPα-AF647 intensity at the contact and outside was quantified and normalized (middle). Linearized fluorescence intensity profiles of the GUVs with the contact highlighted in grey in the confocal image (right). C-E| Single protein controls. Representative confocal images of GUVs comprising Ni-lipid and with B7-H6-AF647 (C), SIRPα-AF647 (D) and CD47-AF488 (E) (left). Fluorescent intensity at the potential contact and outside was quantified and normalized (right). Superplots show individual contacts as small symbols. Large symbols depict the mean of the individual biological replicates. Symbol corresponds to the individual biological replicate (n=3). Standard error of the mean is shown. Scale bar corresponds to 10 μm.

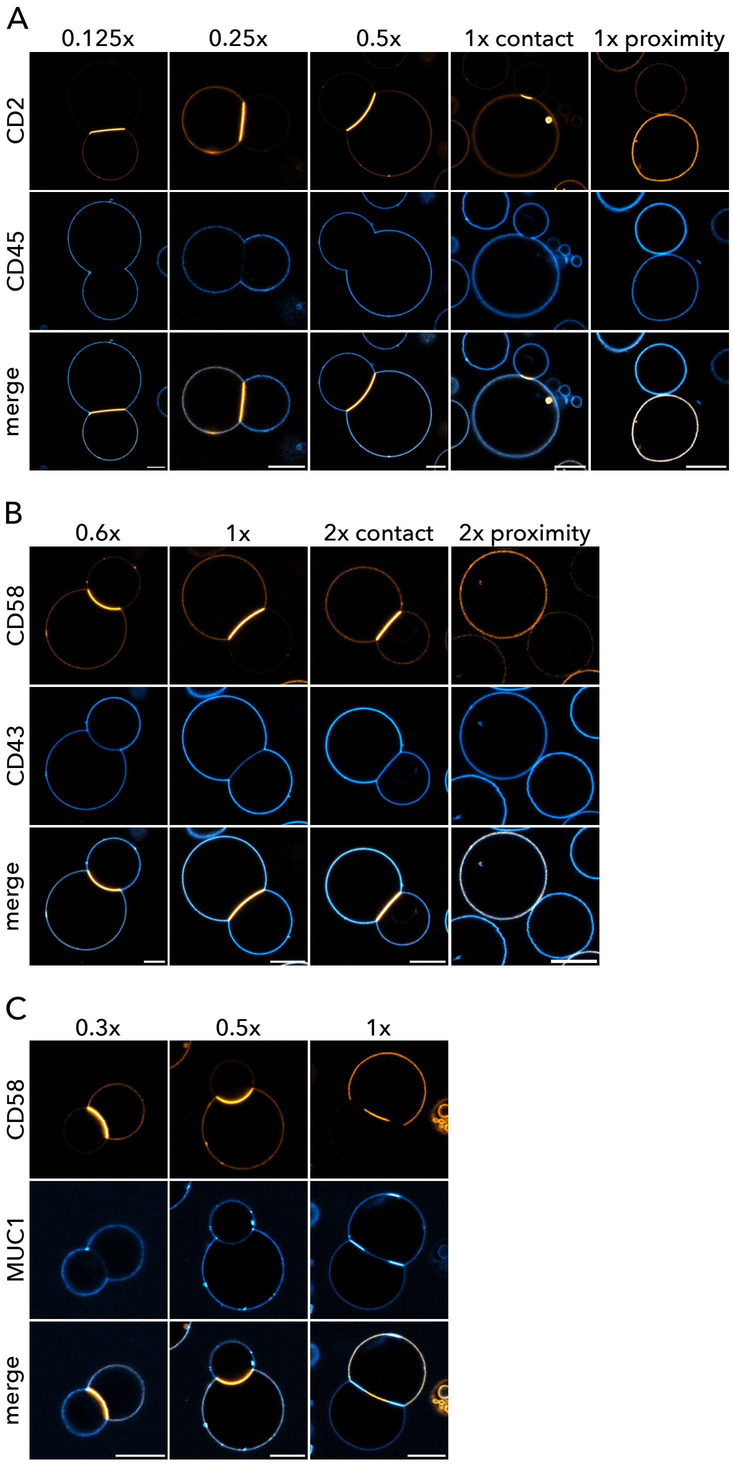


**Supplement Figure 4: Exclusion of CD45, CD43, MUC1 at the CD2-CD58 contact.** A| Representative confocal images of CD45-AF647 at the CD2-AF488 and CD58 (unlabelled) contact. CD45-AF647 was added in increasing amounts (0.125x, 0.25x, 0.5x and 1x). B| Representative confocal images of CD43-AF488 at the CD58-AF647 and CD2 (unlabelled) contact. CD43-AF488 was added in increasing amounts (0.6x, 1x and 2x). C| Representative confocal images of MUC1-AF488 at the CD58-AF647 and CD2 (unlabelled) contact. MUC1-AF488 was added in increasing amounts (0.3x and 0.5x). Scale bar corresponds to 10 μm.

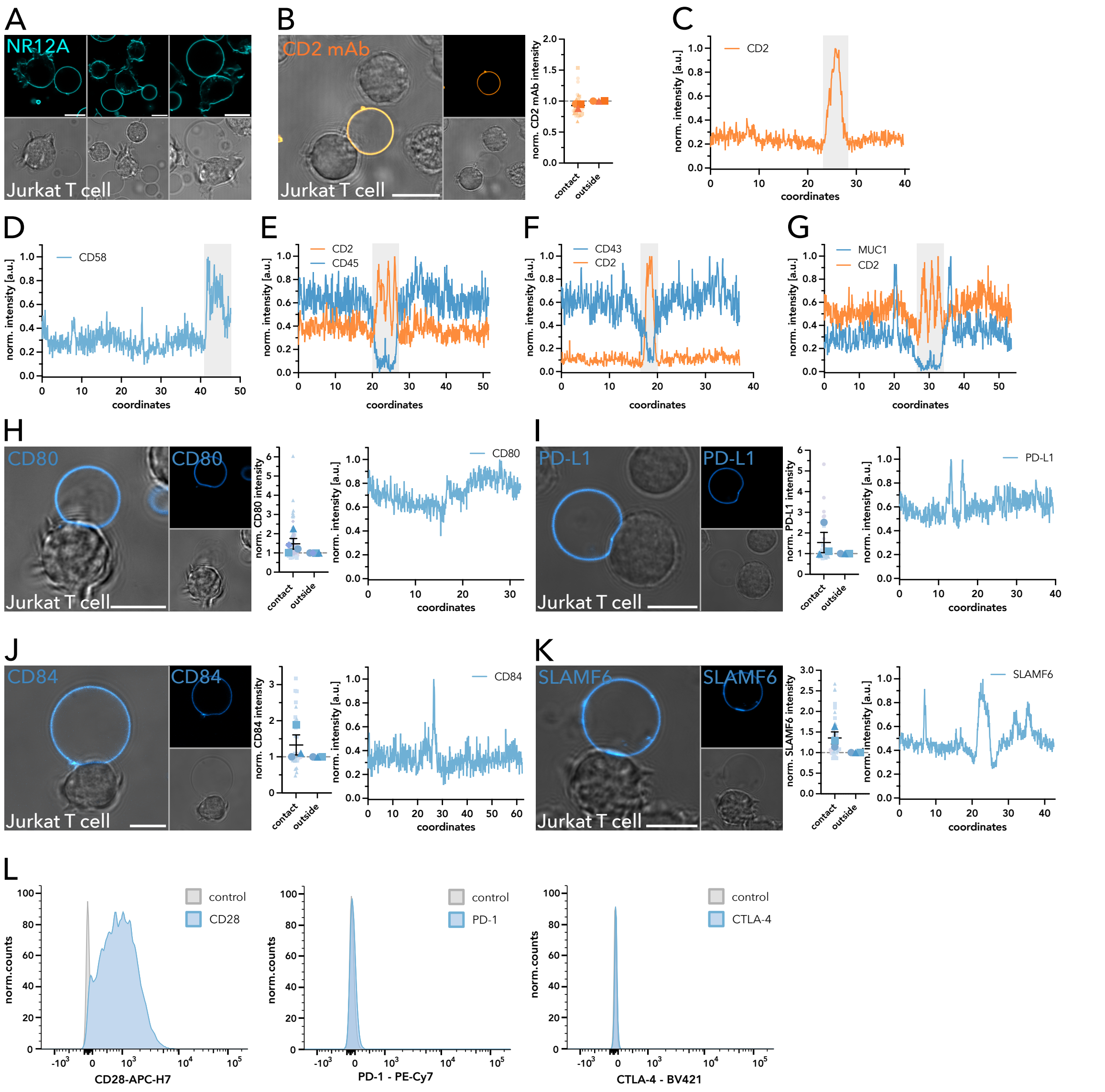


**Supplement Figure 5:** A| Representative confocal images of Jurkat T cells and GUVs comprising Ni-lipid stained with 100 nM NR12A. B| GUVs decorated with unlabelled CD2 and a mAb anti-CD2-AF488 was added to block CD2. Representative confocal images of these GUVs and Jurkat T cells (left). Anti-CD2-AF488 mAb intensity at the potential contact and outside was quantified and normalized (right). C-D| Linearized fluorescence intensity profiles of the GUVs decorated with either CD2-AF488 (C) or CD58-AF647 (D) with the contact highlighted in grey in the confocal images of Figure 2 B-C. E-G| Linearized fluorescence intensity profiles of the GUVs decorated with CD2-AF488 and CD45-AF647 (E); CD2-AF647 and CD43-AF488 (F); or CD2-AF647 and MUC1-AF488 (G) with the contact highlighted in grey in the confocal images of Figure 2 F-H. H-K| Representative confocal images of contact formation between GUVs decorated with CD80 (H), PD-L1 (I), CD84 (J) or SLAMF6 (K) with Jurkat T cells (left). CD80-AF647, PD-L1-AF647, CD84-AF647 and SLAMF6-AF647 intensity at the contact and outside was quantified and normalized (middle). Linearized fluorescence intensity profiles of the GUVs in the confocal image (right). L| Flow cytometry histograms showing the expression of CD28 (left), PD-1 (middle) and CTLA-4 (right) in Jurkat T cells compared to the unstained control. Superplots show individual contacts as small symbols. Large symbols depict the mean of the individual biological replicates. Symbol corresponds to the individual biological replicate (n≥3). Standard error of the mean is shown. Scale bar corresponds to 10 µm.

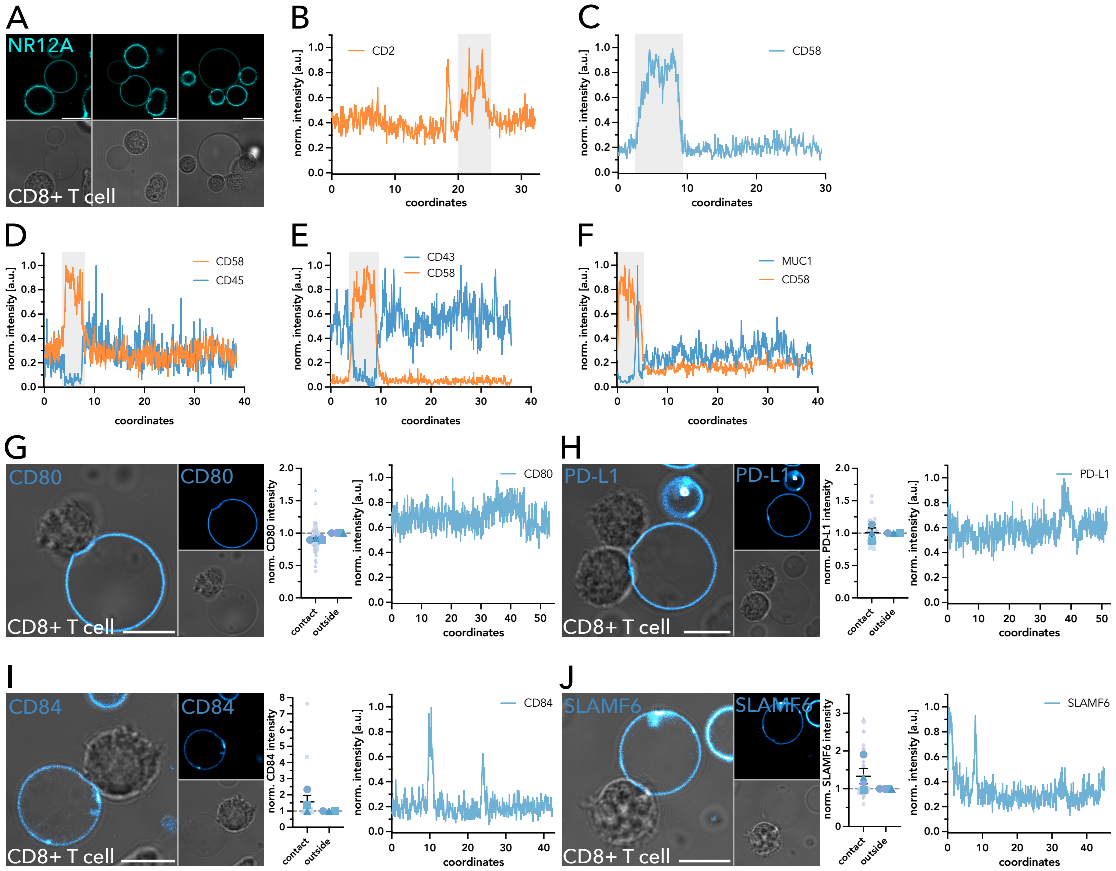


**Supplement Figure 6:** A| Representative confocal images of activated human primary CD8+ T cells and GUVs comprising Ni-lipid stained with 100 nM NR12A. B-C| Linearized fluorescence intensity profiles of the GUVs decorated with CD2-AF488 (B) or CD58-AF647 (C) with the contact highlighted in grey in the confocal images of Figure 2 D-E. D-F| Linearized fluorescence intensity profiles of the GUVs decorated with CD58-AF488 and CD45-AF647 (D); CD58-AF647 and CD43-AF488 (E); or CD58-AF647 and MUC1-AF488 (F) with the contact highlighted in grey in the confocal images of Figure 2 I-K. G-J| Representative confocal images of contact formation between GUVs decorated with CD80 (G), PD-L1 (H), CD84 (I) or SLAMF6 (J) with activated human primary CD8+ T cells (left). CD80-AF647, PD-L1-AF647, CD84-AF647 and SLAMF6-AF647 intensity at the contact and outside was quantified and normalized (middle). Linearized fluorescence intensity profiles of the GUVs in the confocal image (right). Superplots show individual contacts as small symbols. Large symbols depict the mean of the individual biological replicates. Symbol corresponds to the individual biological replicate (n≥3). Standard error of the mean is shown. Scale bar corresponds to 10 μm.

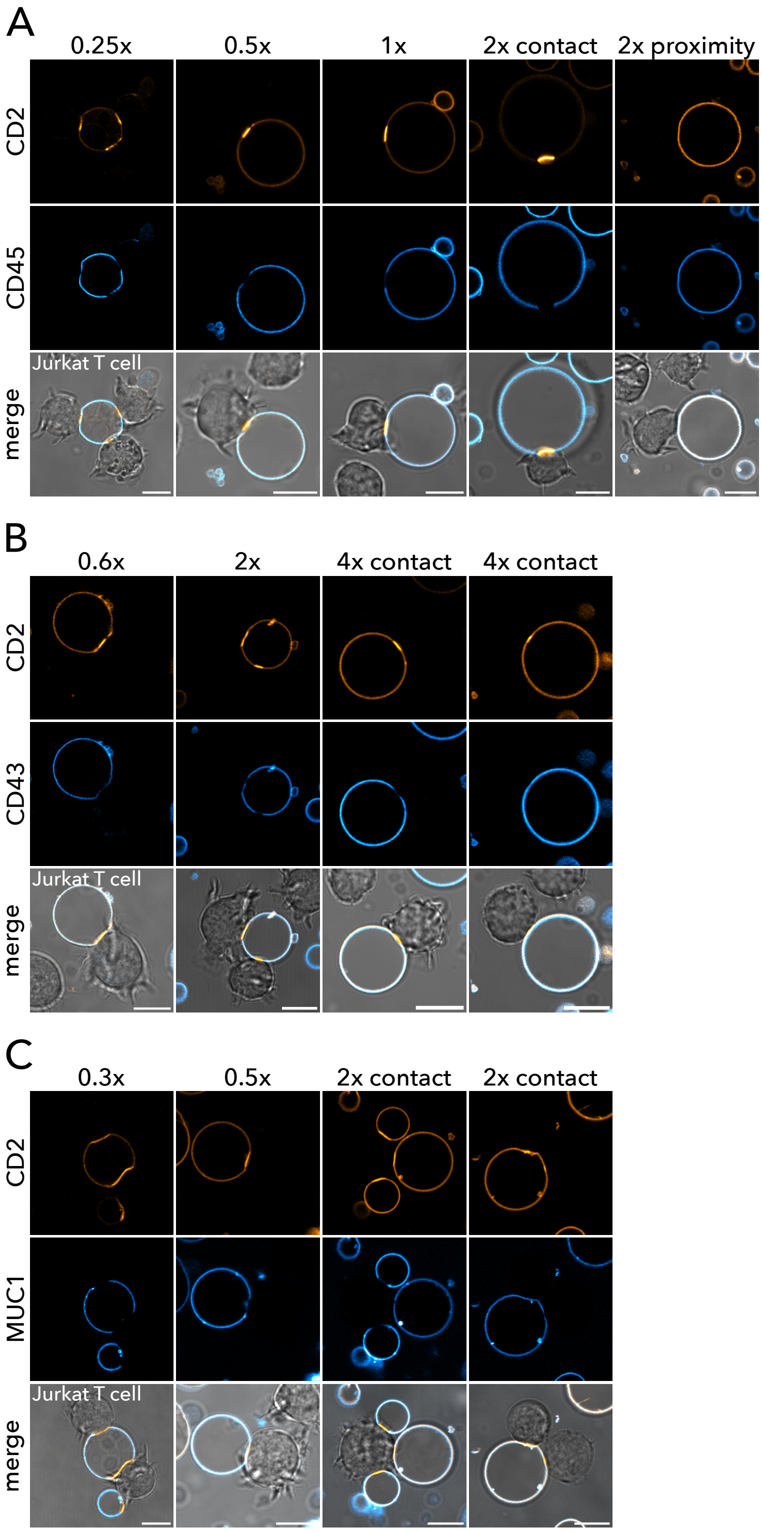

**Supplement Figure 7: Exclusion of CD45, CD43, MUC1 at the CD2-decorated GUV-Jurkat T cell contact.** A| Representative confocal images of contact formation between GUVs decorated CD2-AF488 and CD45-AF647 with Jurkat T cells. CD45-AF647 was added in increasing amounts (0.25x, 0.5x, 1x and 2x). B| Representative confocal images of contact formation between GUVs decorated CD2-AF488 and CD43-AF488 with Jurkat T cells. CD43-AF488 was added in increasing amounts (0.6x, 2x and 4x). C| Representative confocal images of contact formation between GUVs decorated CD2-AF488 and MUC1-AF488 with Jurkat T cells. MUC1-AF488 was added in increasing amounts (0.3x, 0.5 and 2x). Scale bar corresponds to 10 μm.

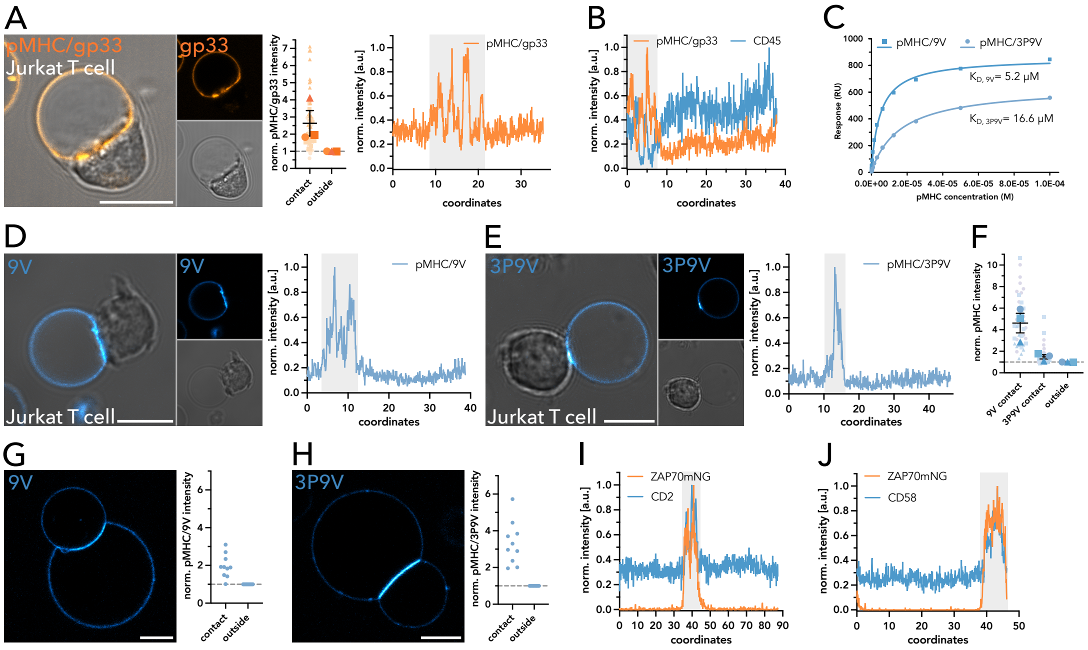


**Supplement Figure 8: T cell activation.**

A| Representative confocal image of contact formation between GUVs decorated with pMHC/gp33 and Jurkat P14 T cells (left). pMHC/gp33-AF488 intensity at the contact and outside was quantified and normalized (middle). Linearized fluorescence intensity profile of GUV decorated with pMHC/gp33-AF488 with the contact highlighted in grey (right). B| Linearized fluorescence intensity profile of GUV decorated with pMHC/gp33-AF488 and CD45-AF647 in the confocal images of Figure 3A. C| BIAcoreTM SPR analysis of the binding of purified 1G4 dsTCR to pMHC/9V and pMHC/3P9V. Response values were plotted against protein concentration. $K_D$ values were obtained from steady-state fitting of equilibrium binding curves from at least ten sample injections. D-E| Representative confocal images of contact formation between GUVs decorated with pMHC/9V (D) or pMHC/3P9V (E) with Jurkat J8 T cells (left). Linearized fluorescence intensity profile of GUVs decorated with pMHC/9V-AF647 or pMHC/3P9V-AF647 with the contact highlighted in grey (right). F| Comparison of quantified and normalized pMHC/9V-AF647 and pMHC/3P9V-AF647 intensities at the contact and outside. G-H| Single protein controls. Representative confocal images of GUVs comprising Ni-lipid and with pMHC/9V (G) or pMHC3P9V-AF647 (H) (left). Fluorescent intensity at the potential contact and outside was quantified and normalized (right). I-J| Linearized fluorescence intensity profiles of the GUVs decorated with CD2-AF647 (I) or CD58-AF647 (J) in the confocal images in Figure 3D with the contact highlighted in grey. Superplots show individual contacts as small symbols. Large symbols depict the mean of the individual biological replicates. Symbol corresponds to the individual biological replicate (n=3). Standard error of the mean is shown. Scale bar corresponds to 10 μm.

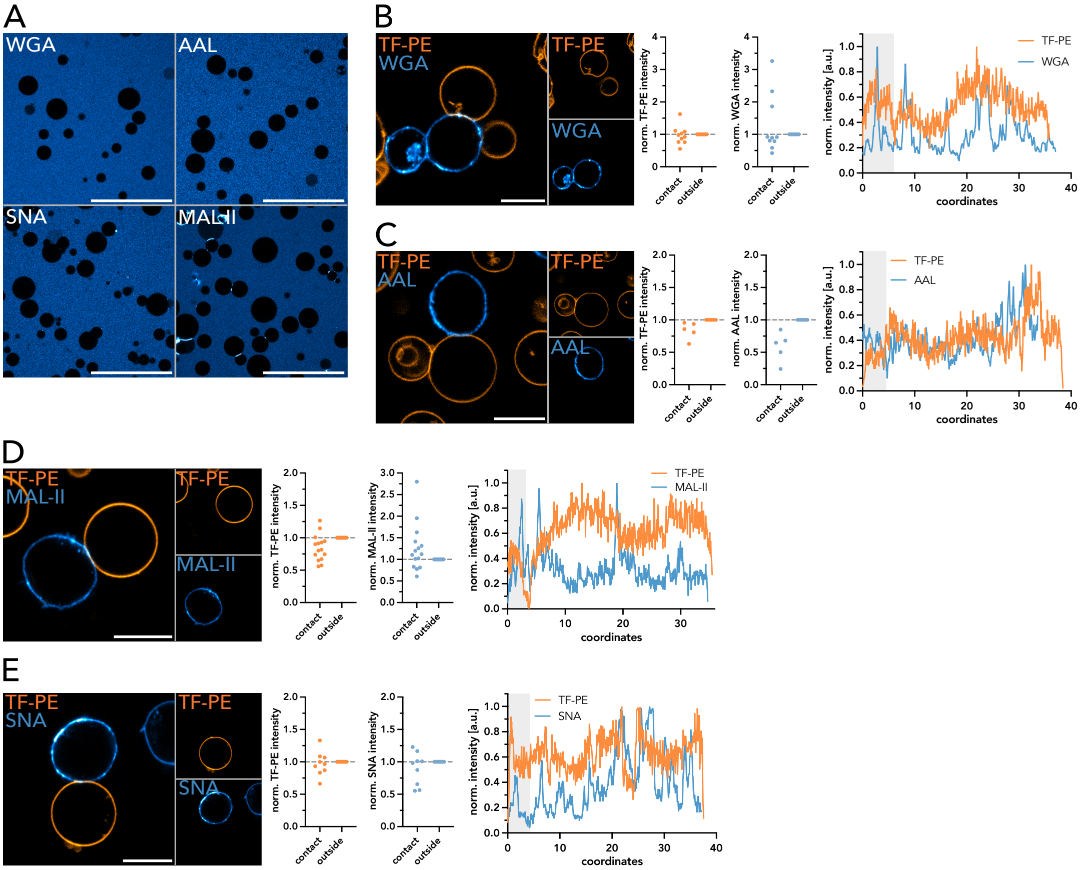


**Supplement Figure 9: Glycocalyx at the T cell GUV contact.** A| Lectin binding controls to no protein-GUVs containing Ni-lipid of WGA-AF647, AAL-AF647, SNA-AF647and MAL-II-AF647. B-E| Representative confocal images of contact formation between no protein-GUVs containing Ni-lipid stained with TF-PE with Jurkat T cells incubated with WGA (B), AAL (C), MAL-II (D) and SNA (E) (left). TF-PE intensity and Lectin intensity at the contact and outside were quantified and normalized (middle). Linearized fluorescence intensity profiles of the GUV and cell stained with the respective lectin in the confocal image with the contact highlighted in grey (right). Superplots show individual contacts as small symbols. Large symbols depict the mean of the individual biological replicates. Symbol corresponds to the individual biological replicate (n=3). Standard error of the mean is shown. Scale bar corresponds to 10μm.
